# A missense mutation in the *Xan-h* gene encoding the Mg-chelatase subunit I leads to a viable pale green line with phenotypic features of potential interest for barley breeding programs

**DOI:** 10.1101/2024.02.24.581867

**Authors:** Andrea Persello, Luca Tadini, Lisa Rotasperti, Federico Ballabio, Andrea Tagliani, Viola Torricella, Peter Jahns, Ahan Dalal, Menachem Moshelion, Carlo Camilloni, Serena Rosignoli, Mats Hansson, Luigi Cattivelli, David Horner, Laura Rossini, Alessandro Tondelli, Silvio Salvi, Paolo Pesaresi

## Abstract

The pale green trait, i.e. reduced chlorophyll content, has been shown to increase the efficiency of photosynthesis and biomass accumulation when photosynthetic microorganisms and tobacco plants are cultivated at high densities. Thus, the *hus1* barley mutant is defective in photosystem antenna biogenesis, and exhibits a 50% reduction in leaf chlorophyll content. Nevertheless, its agronomical performance under standard field conditions is comparable to that of the wild-type. This supports the notion that crops can decrease their investment in antenna proteins and chlorophyll biosynthesis without detrimental effects on productivity. Here, we assess the effects of reducing leaf chlorophyll content in barley by altering the chlorophyll biosynthesis pathway (CBP). To this end, we have isolated and characterised the pale green barley mutant *xan-h.chli-1*, which carries a missense mutation in the *Xan-h* gene for subunit I of Mg-chelatase (*Hv*CHLI), the first enzyme in the CBP. Intriguingly, *xan-h.chli-1* is the only known viable homozygous mutant at the *Xan-h* locus in barley. The Arg298Lys amino-acid substitution in the ATP-binding cleft causes a slight decrease in *Hv*CHLI protein abundance, and a marked reduction in Mg-chelatase activity. Under controlled growth conditions, mutant plants display reduced accumulation of antenna and photosystem core subunits, together with reduced photosystem II yield relative to wild type under moderate illumination, and consistently higher than wild-type levels at high light intensities. Moreover, the reduced content of leaf chlorophyll is associated with a stable reduction in daily transpiration rate, and slight decreases in total biomass accumulation and water-use efficiency. These traits are reminiscent of phenotypic features of wild barley accessions and landraces that thrive under arid climatic conditions. Overall, our findings make the *xan-h.chli-1* allelic variant of potential interest for tailoring barley, and other crop plants, for growth in harsh environments.

## Introduction

Climate change, increasing population growth and scarcity of land undermine the current paradigm of modern agriculture, and our approach to crop production must become more sustainable. While plant architecture and grain yield have been widely explored in modern breeding programs, photosynthetic traits have generally been neglected, and still offer great potential for further crop improvement and adaptation to cope with emerging climatic parameters (Long, Marshall-Colon, and Zhu 2015). The solar Energy Conversion Efficiency (ECE) index, which is defined as the proportion of absorbed radiation that is converted into biomass, relies on whole-canopy photosynthetic efficiency, and overall crop biomass largely depends upon the ECE (Slattery and Ort 2021). This factor is especially relevant, because ECE often falls below half of its theoretical maximum levels in crops (Slattery and Ort 2021). Due to competition for light and nutrients, which are crucial for reproductive success under natural conditions, plants accumulate chlorophylls and thylakoid antenna proteins in large excess with respect to the optimal required for autotrophic growth (Canham et al. 2011). In fact, the photosynthetic machinery saturates at approximately 25% of maximum solar flux in C3 plant canopies, and this represents a major constraint on productivity in these species (Jansson et al. 2010). On the other hand, in anthropic environments, such as cultivated fields characterized by monocultures, competition among individual plants is disadvantageous, and new cultivars with reduced chlorophyll accumulation might become valuable resources. To this end, reduction of leaf chlorophyll content has been suggested to be highly effective in improving light penetration under high-density mass cultivation, and in mitigating high-light-related photo-oxidative damages, with great benefits for biomass yield (Melis 2009). In addition, the reduction of leaf chlorophyll content in crops, *i.e.* the use of pale green phenotypes, enhances light reflectance, which helps to alleviate the effects of heat waves triggered by global climate change (Genesio, Bassi, and Miglietta 2021), and improves the efficiency of water use by reducing canopy temperature (Drewry, Kumar, and Long 2014; Galkin et al. 2018). Furthermore, independent studies have predicted that reductions in chlorophyll content should increase the efficiency of nitrogen use (Walker et al. 2018; Sakowska et al. 2018). Pale green crops can be created by manipulating a plethora of processes, such as the biogenesis and/or accumulation of antenna proteins – also known as the Truncated Light-harvesting Antenna (TLA) strategy – and pigment biosynthesis (for a review, see Cutolo et al., 2023). For instance, increased photosynthetic performance and enhanced plant biomass accumulation were observed upon cultivation at high density under greenhouse conditions of a pale green tobacco line with downregulated expression of *cpSRP43* (Kirst et al., 2018). This nuclear gene codes for the 43-kDa chloroplast-localised signal recognition particle, which is responsible for the delivery of antenna proteins to the thylakoid membranes (Klimyuk et al. 1999), More recently, the barley mutant *happy under the sun 1* (*hus1*), which carries a premature stop codon in the corresponding *HvcpSRP43* gene and is characterised by a 50% reduction in the chlorophyll content of leaves, was shown to accumulate biomass and grains at levels comparable to those observed for the control cultivar Sebastian, when grown under field conditions at standard density. These findings demonstrate that crops can indeed decrease their investment in antenna proteins and chlorophyll biosynthesis significantly, without detrimental effects on productivity (Rotasperti et al. 2022). Conversely, a decrease of about 26% in biomass production was observed in the case of the pale green soybean mutant *MinnGold* under field conditions (Sakowska et al. 2018). Owing to a missense mutation in the nuclear gene encoding the CHLI subunit of the enzyme Mg-protoporphyrin IX chelatase (Mg-chelatase), this mutant synthesizes approximately 80% less chlorophyll than the green control plants. As the first enzyme specific for the chlorophyll biosynthetic pathway, Mg-chelatase is a multimeric complex that is responsible for the insertion of Mg^2+^ into the protoporphyrin IX tetrapyrrole ring. In plants, the enzyme complex consists of three subunits (Masuda 2008), designated CHLI (36–46 kDa), CHLD (60–87 kDa) and CHLH (120–155 kDa), which are also highly conserved in algae and photosynthetic bacteria (Axelsson et al. 2006; Peter and Grimm 2009). In the presence of Mg^2+^ and ATP, the CHLI and CHLD subunits form a double homohexameric ring complex typical of members of the AAA+ [ATPase Associated with various cellular Activities] protein superfamily (Elmlund et al. 2008; Lundqvist et al. 2013), which then interacts with the CHLH subunit responsible for binding protoporphyrin IX and inserting Mg^2+^ to form Mg-protoporphyrin IX (Farmer et al. 2019; Adams et al. 2020; Willows and Beale 1998). The ATP needed for this reaction is hydrolysed by the CHLI subunit which contains the typical AAA+ motifs, such as Walker A and B domains (W-A and W-B), pre-sensor I insert (PS-I), helix 2 insert (H2-insert), arginine finger (ARG-finger) and sensors 1 and 2 (S-1 and S-2) (Lundqvist et al. 2010). The ATP-binding pocket is formed by two neighbouring CHLI subunits via five key interaction motifs (Gao et al. 2020). While most photosynthetic species, including barley, have only one *Hv*CHLI isoform, *Arabidopsis thaliana* has two *CHLI* genes, *AtCHLI1* and *AtCHLI2*, with *AtCHLI1* being more highly expressed than *AtCHLI2* (Huang and Li 2009). Extensive characterization of *chli* mutants has been conducted in various land plants, including Arabidopsis, barley, maize, rice, pea, strawberry and tea (Zhang et al. 2023; Ma et al. 2023; H. Zhang et al. 2006; Wu et al. 2022; Huang and Li 2009; Braumann, Stein, and Hansson 2014). Intriguingly, many forward genetic screens in barley mutant populations have identified chlorophyll-deficient lines with seedling-lethal phenotypes, designated as *Xantha* and *Chlorina* mutants, including *xan-h.38*, *xan-h.56*, *xan-h.57*, *xan-h.clo125*, *xan-h.clo157*, and *xan-h.clo161* (Braumann, Stein, and Hansson 2014; Hansson et al. 1999). These *Xantha* and *Chlorina* mutants carry nonsense and missense mutations, respectively, in the *Xan-h* coding region. Interestingly, heterozygous missense mutations display stronger semidominance than the nonsense mutations (Hansson et al. 2002; Braumann, Stein, and Hansson 2014).

Alongside its key role in chlorophyll biosynthesis, Mg-chelatase plays a vital role in chloroplast-to-nucleus retrograde communication. Thus, inactivation of Mg-chelatase due to a mutation in *CHLH* resulted in the Arabidopsis *gun5* mutant (*genomes uncoupled 5*), which deregulates the expression of the *Light Harvesting Complex B2* gene (*LHCB2*) (Mochizuki et al. 2001) upon inhibition of chloroplast biogenesis. The GUN4 protein (*genomes uncoupled 4*), a regulatory subunit found in oxygenic photosynthetic organisms, which binds to CHLH and stimulates its magnesium chelatase activity, has also been reported to participate in retrograde signalling (Larkin et al. 2003). Similarly, *chld* mutants that are deficient in Mg-chelatase activity show plastid-mediated deregulation of selected nuclear genes (Brzezowski et al. 2016; Huang and Li 2009). With regard to *chli* mutants, it was reported that *A. thaliana cs* and *ch42* and rice *chlorina-9* mutants do not show the *genomes uncoupled* phenotype (Mochizuki et al. 2001; Zhang et al. 2006), whereas both the semi-dominant Arabidopsis mutant *cs215/cs215* and the *Atchli1/Atchli1 Atchli2/Atchli2* double mutant do, since they accumulate higher levels of *Light Harvesting Complex B1* (*LHCB1*) transcripts than the wild type upon impairment of chloroplast activity by norflurazon (NF) treatment (Huang and Li 2009). In barley, lethal mutations in any of the three Mg-chelatase genes cause the *genomes uncoupled* phenotype (Gadjieva et al. 2005).

In this study, we describe the pale green barley mutant line *TM2490*, which was isolated from the TILLMore mutagenized population (Talamè et al. 2008), and is characterised by a single point mutation (Arg-to-Lys at position 298) in the *Xan-h* (*HORVU.MOREX.r3.7HG0738240*) gene. The homozygous mutant plants show reduced leaf chlorophyll content and increased photosynthetic efficiency at high light intensities, and represent the only known viable homozygous *xan-h.chli-1* mutant in barley. In the following, we provide further insights into *Hv*CHLI function and chlorophyll accumulation in barley, and explore the behaviour of the pale green leaf phenotype under drought stress with a view to ultimately exploiting this trait in barley breeding programs.

## Results

### The pale green phenotype of the *TM2490* barley mutant is caused by a missense mutation in *Xan-h*, the single-copy nuclear gene encoding the *Hv*CHLI subunit of Mg-chelatase

The chemically mutagenized TILLMore population (Talamè et al. 2008) was screened for pale green mutants with improved photosynthetic performance under field conditions. Among M_4_ mutant lines, the *TM2490* line was selected based on its reduced chlorophyll content, *i.e.* pale green leaf phenotype and enhanced photosynthetic performance with respect to the control (*cv.* Morex). Under controlled greenhouse conditions, the growth rate and plant architecture of the *TM2490* line were similar to those of the wild-type control. However, amounts of chlorophylls *a* and *b* (Chl*a* + Chl*b*) ranged between 50% of WT levels in the first and second leaves to only 25% in the sixth, i.e. penultimate, leaf, while no major differences in chlorophyll abundance were observed between mutant and control flag leaves (**Fig. 1 A, B**). Similarly, in mutant plants, younger leaves showed a generally increased photosynthetic efficiency of photosystem II (Fv/Fm) relative to the control under both dark-adapted and light-adapted conditions (**Fig. 1 C**), while at later stages no major differences could be observed between mutant and control leaves.

**Figure 1.**
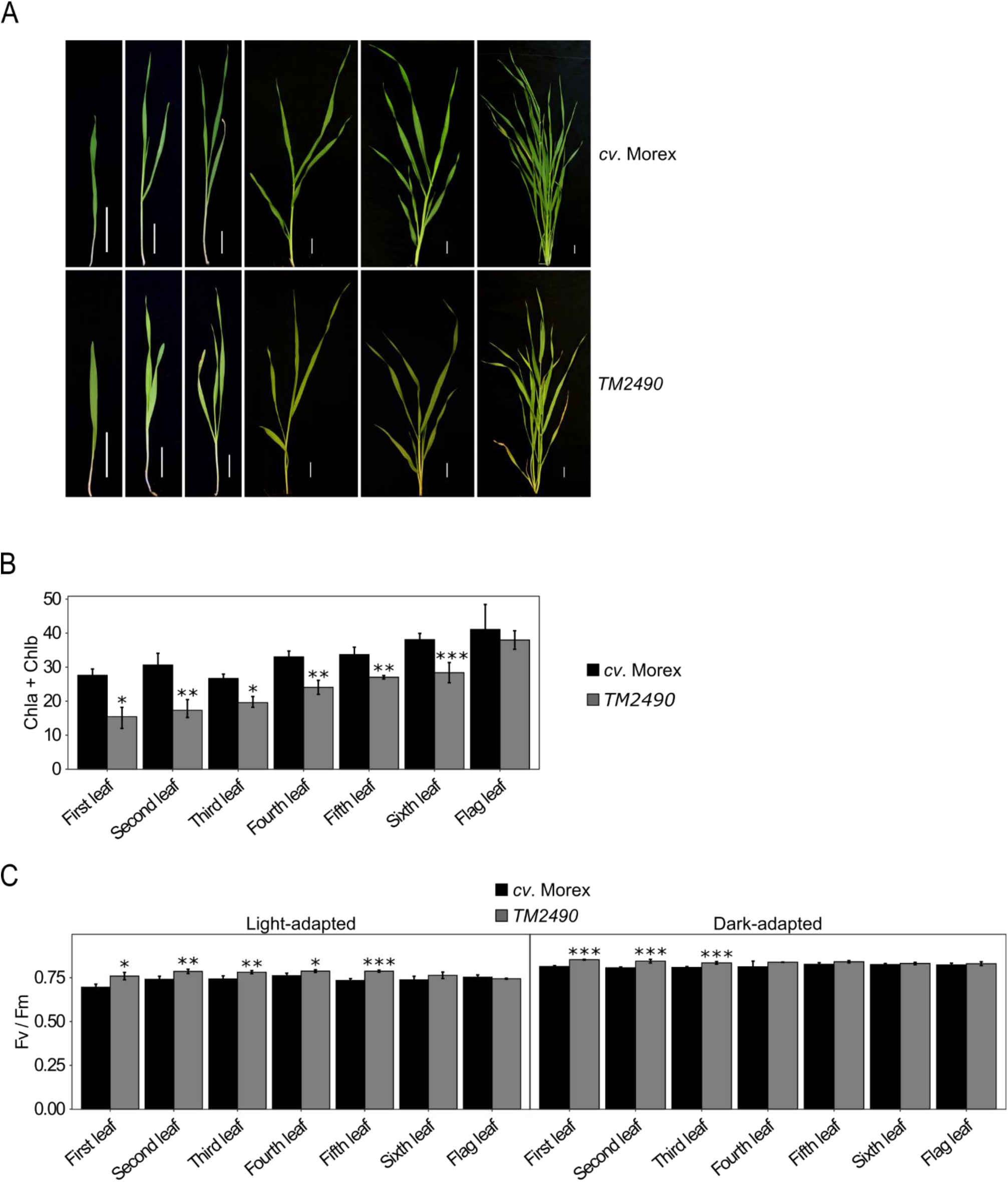
Visible phenotypes of *cv.* Morex (control) and *TM2490* mutant plants grown under greenhouse conditions. (A) Images of *cv*. Morex control plant and the pale green mutant *TM2490* from coleoptile to flag-leaf stage. Scale bar = 2 cm. (B) Measurements of chlorophyll content in *cv.* Morex and *TM2490* leaves (expressed as SPAD units) carried out on eight independent plants at different developmental stages. (C) Leaf photosynthetic performance of dark-adapted and light-adapted plants measured with the Handy PEA fluorometer in eight independent plants. Error bars on the histograms indicate standard deviations and the significance of the observed differences was assessed using Student’s t-test (*** P < 0.001, ** P < 0.01, * P < 0.05).

In order to identify the mutation responsible for the pale green phenotype, a segregating F_2_ population of 565 plants was generated by crossing the *TM2490* line (background *cv.* Morex) with *cv.* Barke. About one-quarter of the total population (131/565) showed the *TM2490*-like phenotype with reduced chlorophyll content and increased Fv/Fm values (WT-like 0.74 ± 0.02 vs *TM2490-like* 0.81 ± 0.02, Student’s t-test <0.001), typical of monogenic recessive inheritance (χ^2^ test 3:1 WT:mut, not significant). Total RNA was then isolated from 100 F2 *TM2490*-like and 100 F2 WT-like plants and bulked in equal ratio to generate two distinct RNA pools. Both RNA pools were subjected to polyA capture and paired-end sequencing, producing approximately 100 million 2 × 150-bp read pairs per pool. Reads were mapped on the reference genome sequence assembly of barley *cv.* Morex (Morex V3; Monat et al., 2019) to identify the allelic variants in each of the two pools. Plotting of the allele frequencies over SNP positions along the barley genome revealed a sharp peak along chromosome 7H, corresponding to a 20-Mb region (from 586.396.977 bp to 606.525.807 bp) in which allelic variants with frequencies higher than 0.5 and peaking at 1.0 in the *TM2490*-like pool were coupled with frequencies lower than 0.5 in the WT-like pool (**Fig. 2 A**). Within this region, the single-copy gene *HORVU.MOREX.r3.7HG0738240*, known as *Xan-h* locus, carried a G-to-A transition at position +1092 from the translation start codon in *TM2490*-like plants (referred to as the *xan-h.chli-1* allele in the following; **Fig. 2 B**), causing an Arg-to-Lys substitution at position 298 (**Fig. S1**). The gene is annotated in the Barlex database as Mg-protoporphyrin IX chelatase subunit I (*Hv*CHLI), a 417-a.a. protein, with a predicted 56-a.a. chloroplast transit peptide (cTP) at the N-terminus, which is essential for the insertion of Mg^2+^ into protoporphyrin IX, the first chlorophyll-specific step of tetrapyrrole biosynthesis in photosynthetic organisms (Kobayashi et al., 2008; Huang and Li, 2009). The protein is highly conserved from photosynthetic bacteria to higher plants as is the Arg residue at position 298 (**Fig. S1**).

**Figure 2.**
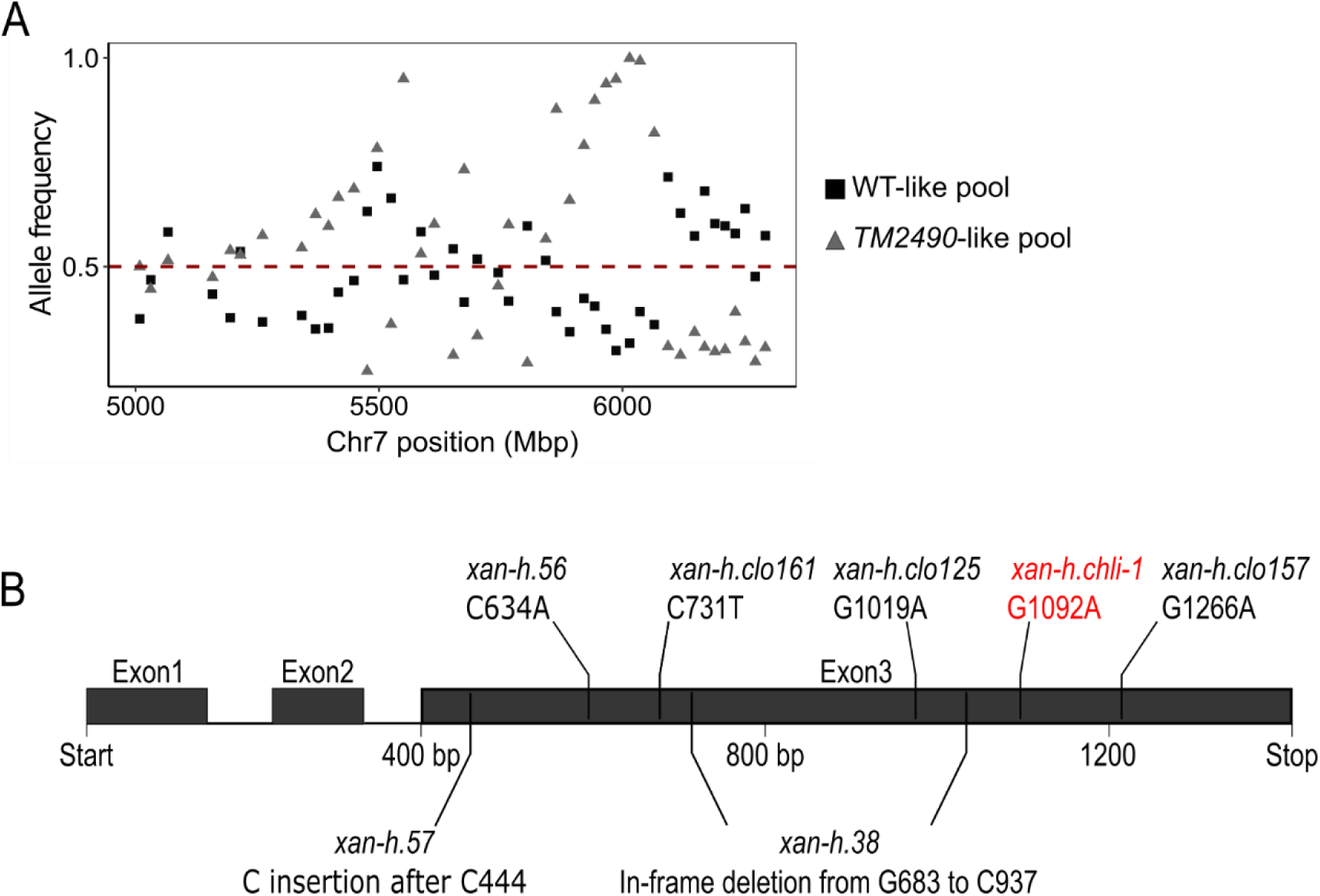
Identification of the *TM2490* locus. (A) Comparison of allele frequency distributions in RNAseq pools obtained from WT-like and *TM2490*-like F2 individuals. Allele frequencies are indicated on the Y axis, genomic coordinates along Chr7 on the X axis. The peak of homozygous alleles in the *TM2490*-like pool corresponds to the 20-Mb candidate region around 6000 Mbp. The red line indicates the threshold allele frequency of 0.5. (B) Schematic representation of the *Xan-h* (*HORVU.MOREX.r3.7HG0738240*) locus, i.e. the single-copy gene chosen as the best candidate for the *TM2490* phenotype. Bars indicate the positions of known lethal mutations within the gene, together with the *TM2490* mutation, here indicated as *xan-h.chli-1*, with the respective SNPs. Boxes represent exons and lines indicate introns.

In order to validate the association between the missense mutation in the *Xan-h* locus and the *TM2490* phenotype, allelism tests were performed, with the aid of the two other known mutant alleles at this locus, *xan-h.clo161* (Hansson et al., 1999) and *xan-h.56* (Braumann, Stein, and Hansson 2014), both of which are recessive chlorotic lethals (see **Fig. 2 B**). To this end, homozygous *TM2490* (*xan-h.chli-1/xan-h.chli-1*) plants were crossed with heterozygous *xan-h.clo161* (*Xan-h*/*xan-h.clo161*) and *xan-h.56* (*Xan-h*/*xan-h.56*) plants, and F_1_ seedlings were phenotypically and genetically analysed at the cotyledon stage. Approximately 50% of F_1_ plants, carrying both *xan-h.chli-1* and either of the *Xan-h* alleles, showed a WT-like photosynthetic and dark-green leaf phenotype, while the biallelic *xan-h.clo161*/*xan-h.chli-1* seedlings were characterised by a dramatic reduction in chlorophyll content, impaired photosynthetic performance (Fv/Fm) and seedling lethality, similar to those of homozygous *xan-h.clo161* mutant seedlings (**Fig. 3 A, C, D**). The pale green phenotype with significant reduction in leaf chlorophyll content was also observed in *xan-h.56*/*xan-h.chli-1* biallelic seedlings, despite showing WT-like photosynthetic performance and the capability to complete the life cycle (**Fig. 3 B, E, F**).

**Figure 3.**
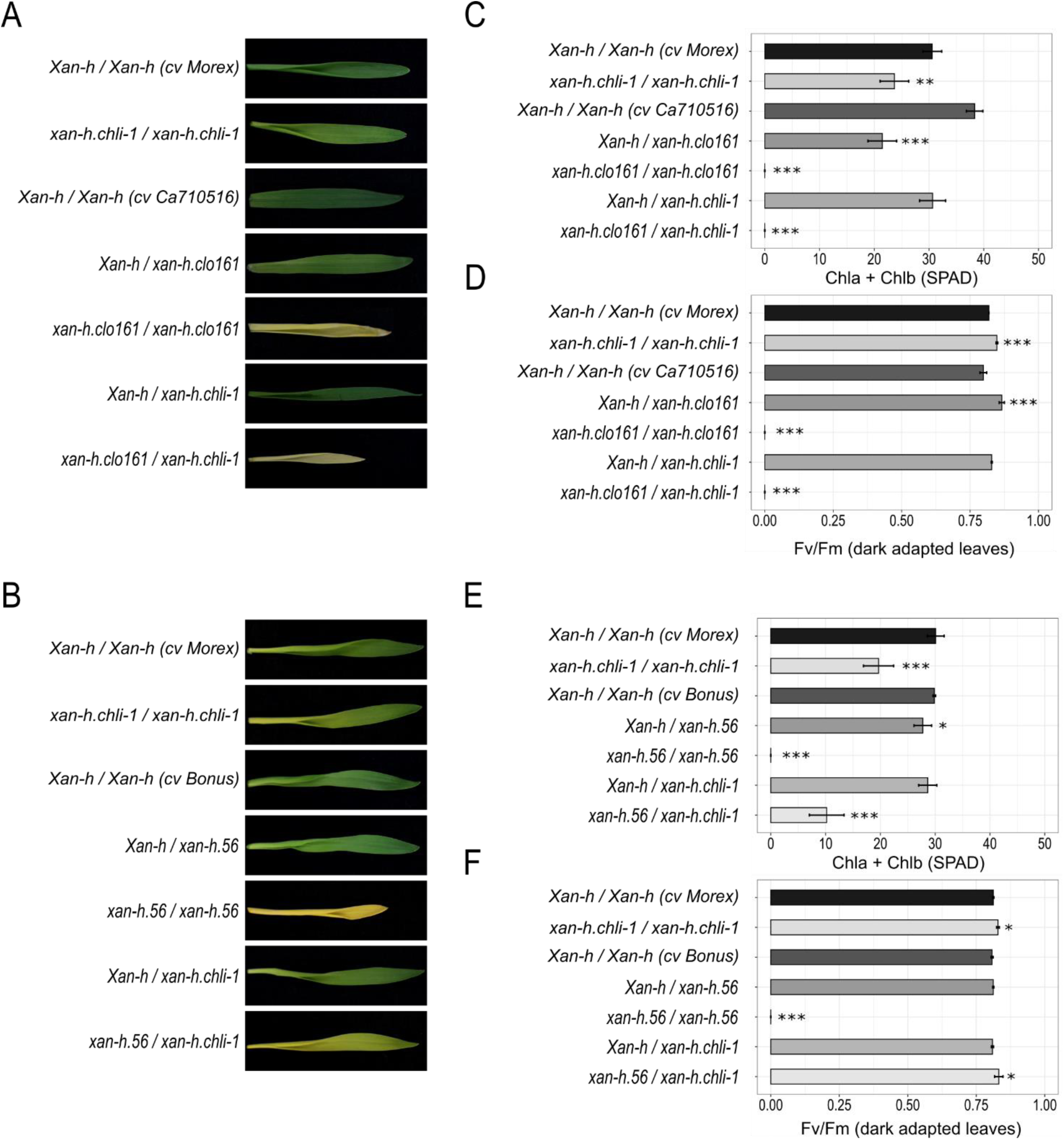
Validation of the *TM2490* locus based on allelism tests. (A and B) First leaves of F1 plants obtained by crossing *TM2490 (xan-h.chli-1/xan-h.chli-1)* with *xan-h.clo161* (genetic background *cv* Ca710516) and *xan-h.56* (genetic background *cv* Bonus), grown for 10 days under controlled greenhouse conditions, together with *cv* Morex, Bonus, Ca710516, *xan-h.chli-1*, *xan-h.56* and *xan-h.clo161*. (C and E) Measurements of total chlorophyll contents expressed in SPAD units. (D and F) Leaf photosynthetic performance of PSII (Fv/Fm) in dark-adapted plants measured using the portable Handy PEA. Error bars on the histograms indicate standard deviations (Student’s t-test; *** P < 0.001, ** P < 0.01, * P < 0.05).

The functional status of the *xan-h.chli-1* mutant allele was further investigated by cloning its coding sequence into a binary vector under the control of the *CaMV35S* promoter and introducing it into the Arabidopsis *Atchli1/Atchli1* knock-out genetic background by Agrobacterium-mediated transformation (Huang and Li, 2009). BLAST analyses indeed revealed that the *Hv*CHLI subunit from *cv.* Morex, encoded by a single-copy gene, shares high homology with the *A. thaliana* proteins *At*CHLI1 (78% identity) and *At*CHLI2 (81% identity) (**Fig. S1**). The *Atchli1/Atchli1+ 35S::Xan-h* line, carrying the WT *HvCHLI* coding sequence from *cv.* Morex and the endogenous Arabidopsis *At*CHLI2 protein, used here as the control, showed a fully complemented phenotype in terms of photosynthetic performance and chlorophyll accumulation in T1 lines and progenies. In contrast, the *35S::xan-h.chli-1* construct only partially complemented the *Atchli1/Atchli1* lethal phenotype, generating viable plant lines that were similar to both barley *TM2490* and Arabidopsis *cs/cs* mutants with respect to photosynthetic performance and total chlorophyll content (**Fig. 4**; Kobayashi et al., 2008). Overall, these data corroborated the hypothesis that the *xan-h.chli-1* mutant allele is responsible for the pale green phenotype and the altered *Hv*CHLI of the homozygous *TM2490* line bearing the R298K amino acid substitution that hampers chlorophyll biosynthesis.

**Figure 4.**
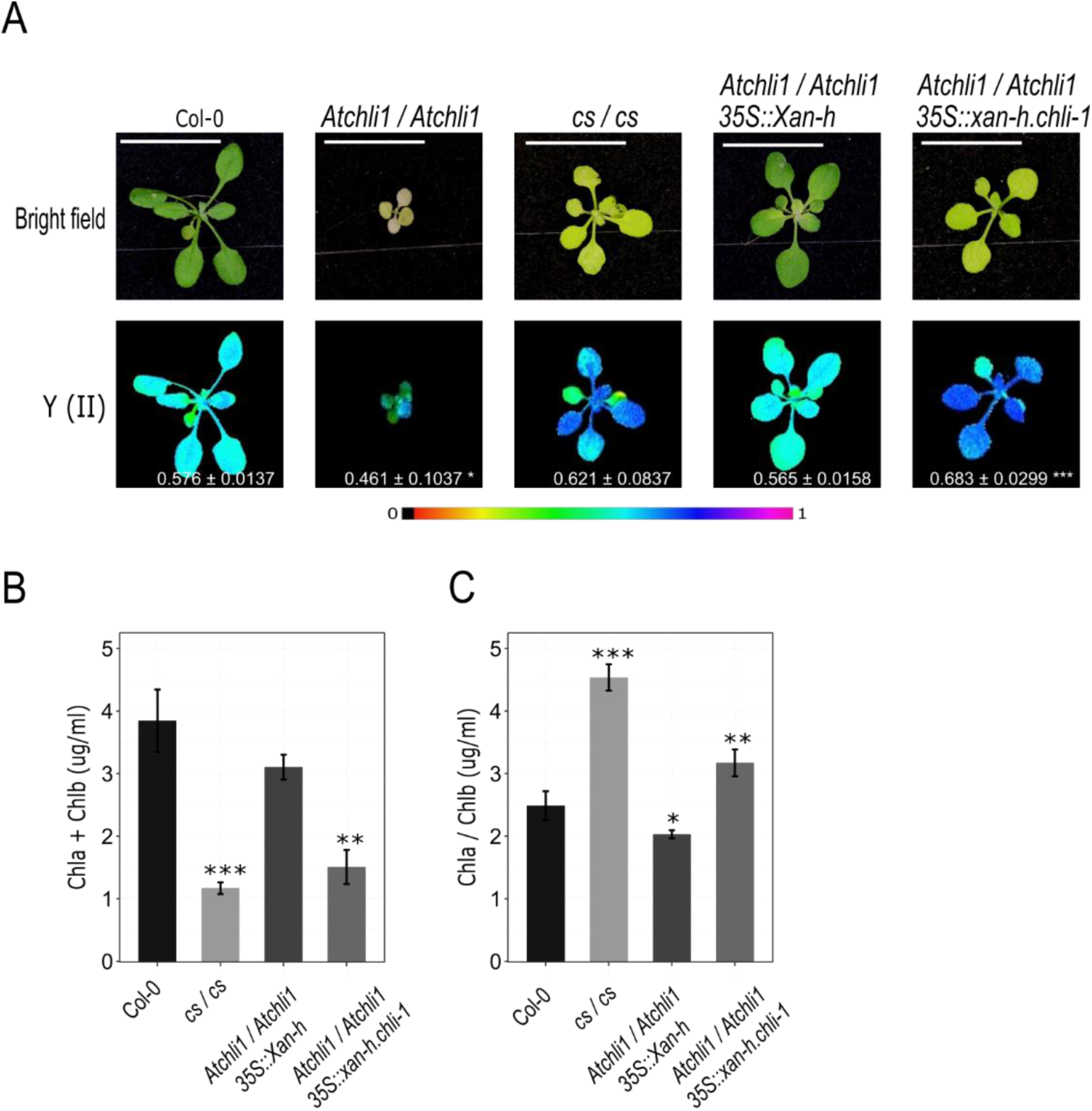
Complementation assays using the *A. thaliana chli1* mutant (*Atchli1/Atchli1*) and the barley WT (*Xan-h*) or the barley mutant allele (*xan-h.chli-1*). (A) Visible phenotype and photosynthetic performance of *Atchli1/Atchli1* rosette leaves compared to the same genotype crossed with either *Xan-h* or *xan-h.chli-1*, and the controls Col-0 and *cs/cs*. Photosynthetic performance was estimated by measuring the Y(II) values of plants that were exposed to light (56 µmol photons m^-2^ s^-1^) for 5 min. The data were recorded with the Imaging-PAM fluorometer and the results are displayed in false colours. The colour scale is shown below the images, where violet corresponds to 1 and black to 0. Scale bar = 1 cm. (B) Chl*a* + Chl*b* content and (C) Chl*a*/Chl*b* ratio determined by spectrophotometry (see Materials and Methods section). Data were analysed with Student’s t-test (*** P < 0.001, ** P < 0.01, * P < 0.05).

### The *xan-h.chli-1* barley mutant shows a reduced chlorophyll content and increased photosynthetic efficiency under high light intensities

To extend the characterization of *TM2490*-related phenotype and minimize any possible influence of other chemically induced mutations in the *TM2490* genome, the mutant was backcrossed with the barley *cv.* Morex, and BC_2_F_2_ plants showing the *TM2490* phenotype (referred to as the *xan-h.chli-1* line in the following) were selected for detailed biochemical and physiological characterization, together with their wild-type-like siblings (referred to as *Xan-h* in the following). In particular, the second leaves of *Xan-h* and *xan-h.chli-1* plants were used for all the analyses reported from here on (**Fig. 5 A**). Quantification of leaf pigments by high-performance liquid chromatography (HPLC) revealed that the total chlorophyll content (Chl*a* + Chl*b*) in *xan-h.chli-1* amounted to about 57% of that in *Xan-h*, while the ratio of Chl*a* to Chl*b* in the mutant (3.97 ± 0.1) was higher than that in the WT (*Xan-h* 3.29 ± 0.1; see also **Table 1**). This difference was due to the reduced accumulation of Chl*a* in *xan-h.chli-1* line (59% of the *Xan-h* level) and an even more marked decrease in Chl*b* (49% of the *Xan-h* level). In addition, the pool of carotenoids associated with photosystem antenna complexes, such as lutein (Lut) and neoxanthin (Nx), showed a marked reduction in the mutant (to around 54% of *Xan-h* levels), while the β-carotene (β-Car) content, found mainly in photosystem cores, was decreased to 65% of *Xan-h* control (**Table 1**), indicating a general alteration of photosystems, albeit more pronounced at the level of antenna proteins. To investigate this aspect further, the second leaves of *Xan-h* and *xan-h.chli-1* plants were exposed to increasing actinic light intensities (0-1287 μmol photons m^−2^s^−1^) and the photosynthetic efficiency was assessed by monitoring the performance of photosystem II (PSII). In dark-adapted leaves, *xan-h.chli-1* showed a higher PSII quantum yield [Y(II)], which declined more rapidly than in *Xan-h* upon moderate light illumination (less than 200 μmol photons m^−2^s^−1^; **Fig. 5 B**). Conversely, the PSII quantum yield of non-regulated energy dissipation [Y(NO)] was markedly higher in *xan-h.chli-1* at low-to-moderate light intensities – implying rather inefficient photochemical energy conversion overall compared to *Xan-h* leaves (**Fig. 5 C**). In addition, upon exposure to 200-1287 μmol photons m^−2^s^−1^ of actinic light, Y(II) values remained consistently higher in *xan-h.chli-1* than in *Xan-h*, possibly because the values for PSII quantum yield attributable to regulated energy dissipation [Y(NPQ)] were consistently lower in *xan-h.chli-1* leaves (**Fig. 5 D**), while Y(NO) levels were identical in mutant and *Xan-h* samples. To investigate further the photosynthetic properties of *xan-h.chli-1* leaves, an identical experimental set-up was used to assess photosystem I (PSI) activity. The quantum yield of PSI [Y(I)] was higher in *xan-h.chli-1* under dark-adapted conditions and dropped to values lower than those seen in *Xan-h*, between 6 and 95 μmol photons m^−2^s^−1^ (**Fig. 5 E**), similarly to the Y(II) trend, and most probably because of less efficient energy transfer from the antenna to the PSI reaction centre. As a matter of fact, the Y(NA) parameter, i.e. the quantum yield of non-photochemical energy dissipation in PSI due to acceptor-side limitation (**Fig. 5 F**), was much higher in *xan-h.chli-1* under low to moderate light conditions, while it decreased to *Xan-h* values at higher light levels, as soon as the photosynthesis control was engaged (Colombo et al. 2016). Similarly, the lower values of Y(ND), *i.e.* the non-photochemical PSI quantum yield due to donor-side-limited heat dissipation (**Fig. 5 G**), at higher light intensities confirmed the greater efficiency of electron transport from PSII to PSI in *xan-h.chli-1* leaves. Overall, our findings highlight the low photosynthetic efficiency of *xan-h.chli-1* under low-to-moderate actinic light intensities, although this parameter rises at higher intensities, most probably as a consequence of the reduced chlorophyll content and light absorption capacity of the pale green *xan-h.chli-1* leaves.

**Figure 5.**
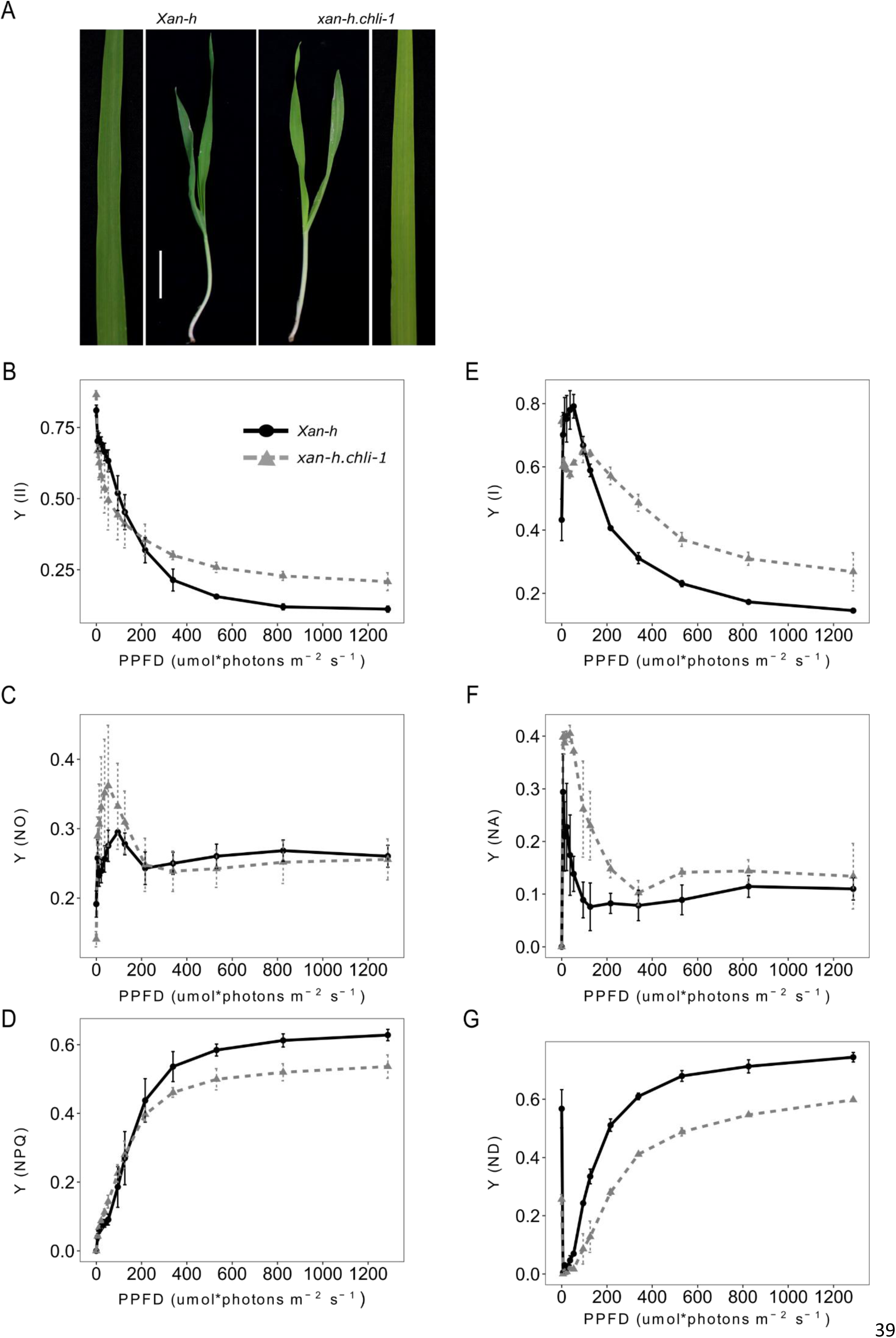
Representative phenotypes of *Xan-h* and *xan-h.chli-1* plants at the second-leaf stage following growth under greenhouse conditions. (A) *Xan-h* and *xan-h.chli-1* barley leaves were harvested 14 days after germination. Note that, in terms of leaf pigment content and photosynthetic performance, *xan-h.chli-1* plants (BC_2_F_2_ generation) were identical to *TM2490* plants at the M4 generation. Scale bar = 2 cm. Analyses of photosynthetic parameters were performed using the Dual-PAM 100 fluorometer. (B) The effective quantum yield of PSII [Y(II)], and quantum yields of (C) non-regulated energy dissipation [Y(NO)] and (D) regulated energy dissipation of PSII [Y(NPQ)]. Measurements used to monitor PSII performance were carried out at increasing light intensities (from dark to 1287 μmol photons m^−2^ s^−1^; 3-min exposure to each light intensity). Concomitantly, the effective quantum yield of PSI [Y(I)] (E), and the quantum yields of non-photochemical energy dissipation in PSI owing to acceptor-side limitation [Y(NA)] (F), and donor-side-limited heat dissipation [Y(ND)] (G), were determined. Curves show average values of three biological replicates, while bars indicate standard deviations. PPFD = photosynthetic photon flux density.

**Table 1.** HPLC analysis of second-leaf pigment content in *Xan-h* and *xan-h.chli-1*. The pigment content was normalized to leaf fresh weight (FW) and is reported as pmol per mg of FW. Nx = neoxanthin; Lut = lutein; Chl = chlorophyll; β-Car = β-carotene; VAZ = violaxanthin + antheraxanthin + zeaxanthin. The significance of the observed differences was evaluated with Student’s t-test (*** P < 0.001, ** P < 0.01, * P < 0.05, ns = not significant).

| | Nx | Lut | Chlb | Chla | β-Car | VAZ | Chla + Chlb | Chla / Chlb |
| --- | --- | --- | --- | --- | --- | --- | --- | --- |
| *Xan-h* | 62±11 | 189±29 | 478±101 | 1563±302 | 182±44 | 120±20 | 2041±402 | 3.29±0.1 |
| *xan-h.chli-1* | 34±9 | 103±30 | 235±52 | 930±203 | 117±30 | 104±25 | 1164±255 | 3.97±0.1 |
| **T-test** | ** | ** | *** | ** | * | ns | ** | *** |

The functional status of the photosynthetic machinery was also analysed at the biochemical level by monitoring the protein composition of the thylakoid electron transport machinery by means of immunoblot analysis. In agreement with the pigment accumulation profile (**Table 1**), the bands corresponding to PSII and PSI light-harvesting proteins (Lhcb and Lhca antenna proteins, respectively) observed in the 20-to 25-kDa range of the SDS-PA gel after staining with Coomassie-Brilliant Blue were markedly decreased (**Fig. 6 A**). Accordingly, immunoblot analyses with antibodies specific for Lhca3, Lhcb1, Lhcb2 and Lhcb3 confirmed a general reduction (of at least 60%) in antenna proteins in *xan-h.chli-1* thylakoids, while lesser declines (of 20-40%) were observed for Lhca1, Lhca2 and Lhcb4. Only in the case of Lhcb5 were the levels attained identical between mutant and *Xan-h* samples (**Fig. 6 A**). Moreover, in *xan-h.chli-1* thylakoid samples the levels of PSII core subunits (D1 and CP43) were reduced by around 30% and 50%, respectively, similar to what was observed for PsbS and two subunits of the Oxygen-Evolving Complex (OEC), PsbO and PsbR. Conversely, the PsbQ subunit of OEC showed a much more drastic reduction, accumulating to only around 14% of its level in *Xan-h*. Similarly to PSII, the PSI core subunits, PsaA and PsaD, accumulated in *xan-h.chli-1* thylakoids to lower levels than in *Xan-h*, while no major differences were observed in the accumulation of the cytochrome *f* subunit (PetA) or the plastocyanin electron carrier (PetE). Finally, the impact of the decreased chlorophyll and thylakoid protein contents on the chloroplast ultrastructure in the mutant was investigated by Transmission Electron Microscopy (TEM; **Fig. 6 B**). TEM analyses, performed on growth-light-adapted plants, showed reduced accumulation of starch granules in *xan-h.chli-1* chloroplasts when compared to *Xan-h*, while no major alteration in the organization of grana and stroma lamellae was observed.

**Figure 6.**
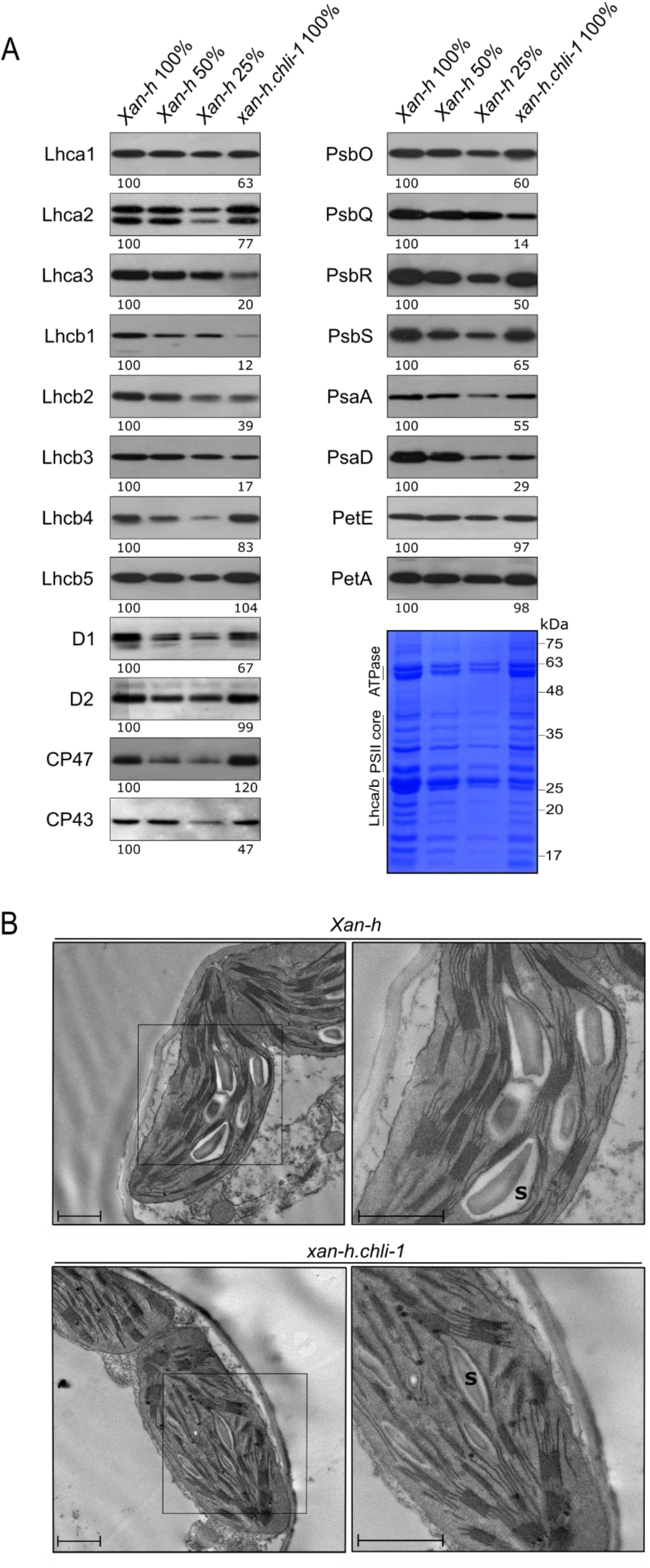
Biochemical and ultrastructural characterization of thylakoid membranes from *Xan-h* and *xan-h.chli-1*. (A) Immunoblot analyses of thylakoid protein extracts from *Xan-h* and *xan-h.chli-1* leaf material, normalized with respect to fresh weight and probed with antibodies specific for subunits of thylakoid protein complexes. For relative quantification, 50% and 25% dilutions of *Xan-h* protein extracts were also loaded. One filter (representative of three biological replicates) is shown for each immunoblot. An SDS-PA gel stained with Coomassie Brilliant Blue (CBB) is shown as loading control. (B) TEM micrographs depict chloroplast ultrastructure in *Xan-h* (upper panels) and *xan-h.chli-1* (lower panels) samples. S = starch granule; Scale bar = 1µm.

### The *xan-h.chli-1* mutation impairs Mg-chelatase activity

Since the *xan-h.chli-1* mutation results in the replacement of the highly conserved Arg residue at position 298 by Lys (**Fig. S1**), the accumulation and activity of Mg-chelatase enzyme was quantified. Immunoblot analyses revealed that the *Hv*CHLH subunit accumulated to WT levels, while *Hv*CHLI and *Hv*CHLD were slightly reduced in *xan-h.chli-1* leaves (**Fig. 7 A**). On the other hand, the abundance of the regulatory subunit *Hv*GUN4 was almost twice as high in *xan-h.chli-1* as it was in *Xan-h*, possibly as a compensatory response to the decline in Mg-chelatase activity owing to partial impairment of *Hv*CHLI. In order to test whether the point mutation identified in the *xan-h.chli-1* allele affects its homodimerization, its coding sequence (devoid of the cTP-coding region) was tested for homodimer formation in a yeast two-hybrid assay, together with the variants *xan-h.clo125*, *xan-h.clo157* and *xan-h.clo161* and compared with the ability of *Xan-h* from *cv.* Morex to homodimerize as a control. As shown in **Fig. 7 B**, colonies expressing *Xan-h* and *xan-h.chli-1* were able to grow on selective media, suggesting that the R298K missense mutation does not impair homodimer formation. In contrast to this result, colonies expressing *xan-h.clo125*, *xan-h.clo157* and *xan-h.clo161* variants were unable to grow on selective media, indicating that lethal mutations hamper the ability to form CHLI homodimers (**Fig. 7 B, Fig. S2**). Moreover, stromal protein extracts from etiolated *Xan-h* and *xan-h.chli-1* seedlings were used to test the activity of the Mg-chelatase oligo-enzyme by measuring its ability to convert deuteroporphyrin IX into Mg-deuteroporphyrin IX *in vitro*. As expected, a marked reduction in Mg-chelatase activity was observed in the *xan-h.chli-1* protein relative to *Xan-h* samples and the mock control **(Figure 7 C)**.

**Figure 7.**
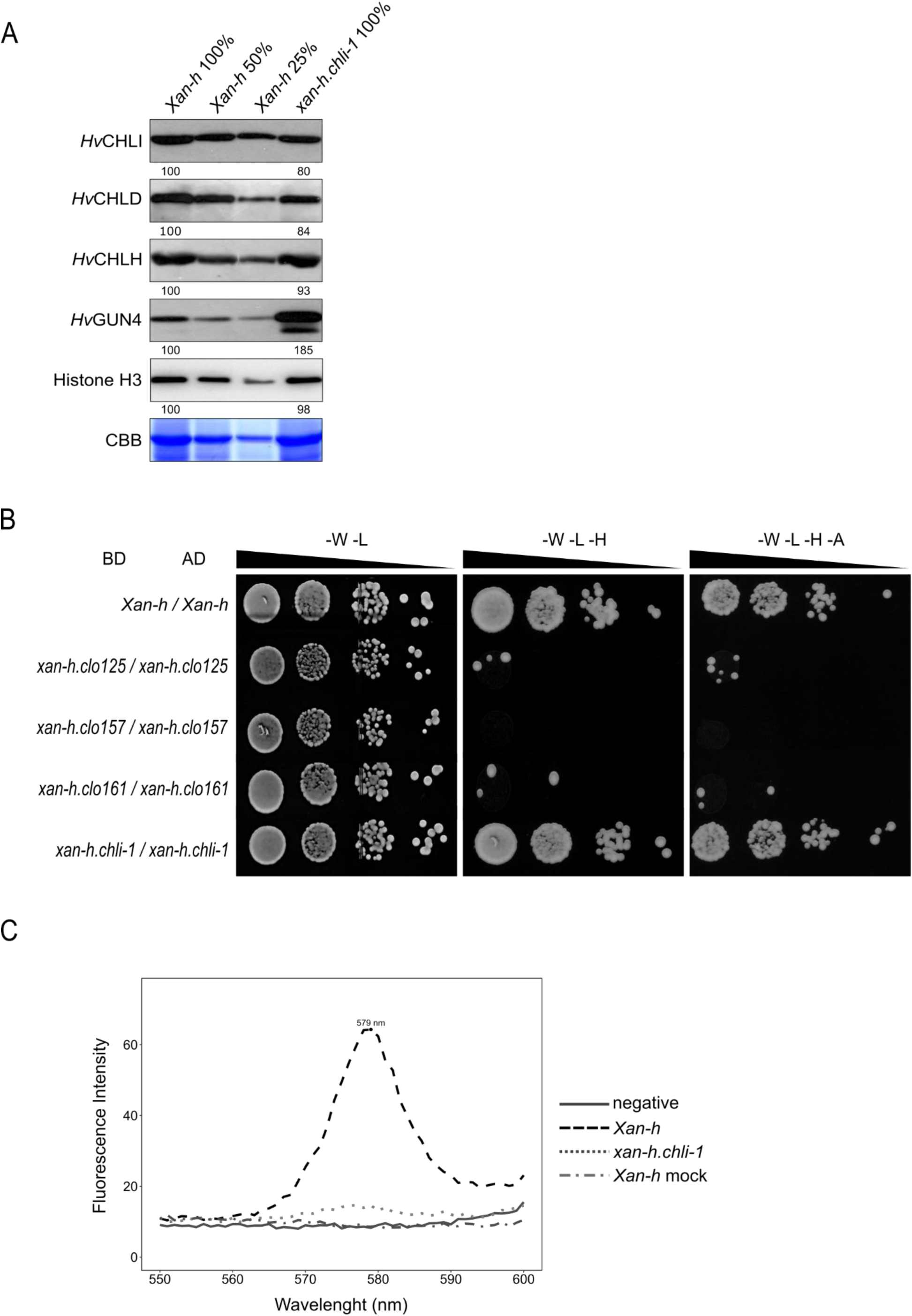
Effects of the *xan-h.chli-1* mutation on the accumulation, assembly and activity of the Mg-Chelatase complex. (A) Immunoblot analyses of total protein extracts (normalized to leaf fresh weight) from *Xan-h* and *xan-h.chli-1* plants with antibodies specific for *Hv*CHLI, *Hv*CHLD, *Hv*CHLH and *Hv*GUN4, respectively. A CBB-stained gel corresponding to the RbcL region, and an immunoblot showing the histone H3 protein are shown as controls for equal loading. For protein quantification, 50% and 25% dilutions of *Xan-h* protein extracts were also loaded. One representative of three biological replicates is shown for each immunoblot. (B) Yeast two-hybrid interaction assay**s** were performed on *Xan-h* and the mutant allelic variants *xan-h.chli-1*, *xan-h.clo125*, *xan-h.clo157* and *xan-h*.*clo161* in order to test their ability to self-interact (homodimerization). As highlighted by their growth on selective media (-W-L-H and –W-L-H-A), only the colonies expressing the wild-type *Xan-h* and its mutant *xan-h.chli-1* alleles were able to self-associate. BD, GAL4 DNA-binding domain, AD, GAL4 activation domain, –W –L, dropout medium devoid of Trp and Leu (permissive medium); –W –L –H, lacking Trp, Leu and His (selective medium), and –W –L –H –A, lacking Trp, Leu, His and Ade (selective medium). serial dilutions were prepared for each strain. (C) *In vitro* assay of Mg-chelatase activity in etiolated leaf extracts from *Xan-h* and *xan-h.chli-1*. The fluorescence emission of the Mg-chelatase product Mg-deuteroporphyrin was recorded from 550 to 600 nm using an excitation wavelength of 408 nm. Protein extracts were normalized to total protein content. One representative chart (of three biological replicates) is shown.

### The R298K substitution in *Hv*CHLI may hamper its interaction with ATP

To analyse the consequences of the R298K substitution in a structural context, we modelled the configuration of the *Hv*CHLI subunit, as described in the Materials and Methods section. This ATPase motor is found in all kingdoms of living organisms and shares a common core structure, although it is involved in different activities and performs diverse functions (Ogura and Wilkinson 2001). To date, the structure has been solved for several organisms other than barley (Gao et al. 2020; Cha et al. 2010; Miller et al. 2014). In general, the *Hv*CHLI ATPase subunit assembles as a closed ring and its quaternary structure results from the association of six identical monomers (Hansson et al. 2002; Lundqvist et al. 2010; **Fig. 8 A**). In the reconstructed model, R298 protrudes towards the ATP-binding cleft at the interface between two monomers (**Fig 8 B**). This position is consistent with the X-ray structure of the *Synechocystis sp. PCC 6803 substr. Kazusa* CHLI subunit (PDB ID 6L8D; Gao et al., 2020), in which R298 corresponds to R233. A similar homology was also observed for other relevant, highly conserved residues, such as R356 (R291 in *Synechocystis*) and D274 (D209 in *Synechocystis*) (**Fig 8 B**). These two residues, whose replacements lead to lethal mutations in *xan-h.clo125* (D274N) and *xan-h.clo157* (R356K), are also located at the ATP-binding pocket. In particular, D274 and R356 belong to different alpha-helices of the same chain and interact with each other (**Fig. 8 C** and **Fig S3A**), establishing an intra-monomer hydrogen bond network that includes R393. The disruption of this interaction could affect the folding of the monomer and thus the formation of CHLI dimer (**Fig. 7 B** and **Fig. 8 C**). On the other hand, the R298K substitution does not lead to either the loss or formation of significant intra-or inter-monomer contacts (**Fig. 8 C**). To further investigate the possible role of R298 in the context of the ATP binding site, the structural analysis was extended to AAA+ proteins that are not related to photosynthesis. In particular, the hexameric structure of the chaperone Heat Shock Locus U [HSLU, PDB ID: 1DO0; (Bochtler et al. 2000)] from *Escherichia coli*, which has been resolved by X-ray analysis with its magnesium ion and ATP in the binding cleft, was utilised to this purpose. The high degree of homology between the ATP-binding domain of HSLU and the barley *Hv*CHLI model enabled us to infer distances between the magnesium ion and the conserved residues in both structures (**Fig. S3 B**). With this information, the magnesium ion was positioned in the binding cleft of the model, and the distances were used to constrain the ATP docking mode (**Fig. 8 B, C**). The results show that R356, whose homologue in *Synechocystis* has been described as part of the S-2 motif, can establish direct hydrogen-bond interactions with ATP (**Fig. 8 C**, left panel), as does the Arg finger R275 (R210 in *Synechocystis*). Hydrogen bonds could also be established with the ATP molecule by the side-chain of R298 (**Fig. 8 C**, left panel). This interaction could be perturbed or prevented by the R298K missense mutation, owing to the shorter side-chain and the presence of only one amino group in lysine (**Fig. 8 C**, right panel). Overall, these observations suggest that R298K missense mutation might compromise the interaction of *Hv*CHLI dimers with ATP.

**Figure 8.**
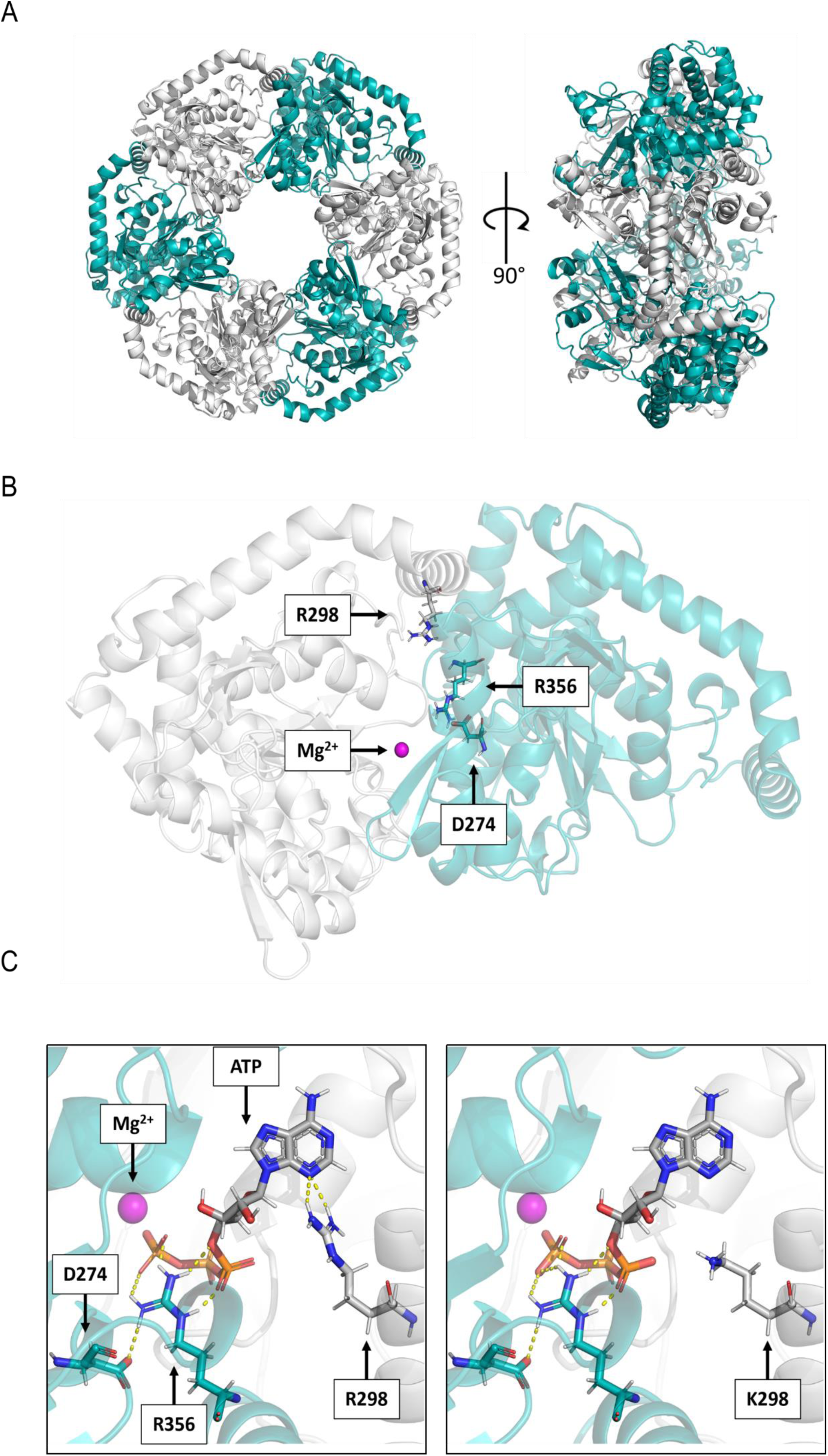
Effect of the *xan-h.chli-1* mutation on the *Hv*CHLI hexamer structure. (A) Model of the AAA+ ATPase subunit of the barley Mg-chelatase enzyme. The adjacent monomers are coloured in white and cyan. Left panel: Frontal view of the homo-hexameric ring shown in cartoon representation; Right panel: Side view of the ring in cartoon representation. (B) Frontal view of a single dimer. The two monomers are represented in transparent cartoon and coloured in white (chain A) and cyan (chain B), respectively. The constituent atoms of R298 in chain A are depicted in light grey (C atoms), blue (N), red (O), and white (H). D274 and R356 (S-2) of chain B are shown in cyan (C), blue (N), red (O), white (H) and represented as solid sticks. R298 in chain B, D274 and R356 from chain A are not highlighted. Mg^2+^ is shown as a magenta sphere. (C) ATP binding cleft with Mg^2+^ and docked ATP. The same colour scheme is used for D274 and R356 of chain B, with ATP shown in dark grey and P atoms in orange H-bonds are represented as yellow dashed lines.

### The reduced Mg-chelatase activity in *xan-h.chli-1* plants does not affect plastid-to-nucleus retrograde signaling or the expression of photosynthesis-associated nuclear genes

Since Mg-chelatase activity is markedly reduced in *xan-h.chli-1* chloroplast, we investigated the possibility that Arabidopsis and barley plants carrying the *xan-h.chli-1* allele might show the *genomes uncoupled* (*gun*) phenotype. To do so, barley *Xan-h*, *xan-h.chli-1* and *xan-h.56* seedlings were grown on MS medium in the presence or absence of norflurazon (NF), and levels of *Lhcb3* and *Rbcs* transcripts were determined. As shown in **Figure S4 A**, *Xan-h* and *xan-h.chli-1* seedlings were able to down-regulate the expression of *Lhcb3* and *Rbcs* genes in the presence of NF, indicating that *xan-h.chli-1* mutant does not display the *gun* phenotype, unlike the lethal *xan-h.56* allele used here as a positive control (Gadjieva et al. 2005). Similarly, Arabidopsis lines carrying either the *Xan-h* or the mutant *xan-h.chli-1* allele from barley under the control of *35SCaMV* promoter markedly reduced the expression of *Lhcb3* and *Rbcs* genes in the presence of NF, like Col-0 and the *cs* mutant. As expected, the *gun5* mutant failed to repress the expression of *Lhcb3* and *Rbcs* genes, further supporting the notion that the *xan-h.chli-1* mutation does not affect plastid-to-nucleus communication (**Fig. S4 B**). To further investigate the impact of the *xan-h.chli-1* mutant allele on leaf gene expression, a transcriptomic analysis was performed on *Xan-h* and *xan-h.chli-1* second leaves obtained from plants grown under greenhouse conditions. Principal component analysis (PCA) revealed that the four transcriptome replicates of each genotype clustered together in two clearly separated groups (**Fig. 9 A**). Moreover, differentially expressed genes (DEGs) were identified by filtering for the log-fold-change (logFC) and the adjusted p-value (padj), which resulted in the identification of 432 up-regulated and 335 down-regulated genes in *xan-h.chli-1* relative to *Xan-h* (**Fig. 9 B; Supplementary data**). The relatively small number of DEGs agrees with the moderate distance between the two PCA clusters. This observation, together with the fact that Biological Process Gene Ontology (GO) term analysis resulted in no significant GO term enrichment, indicates that the *xan-h.chli-1* mutant allele does not cause major changes at the transcriptional level with respect to its *Xan-h* counterpart. SUBA5 location prediction was applied to the Arabidopsis homologs of the up and down-regulated genes. Among the up-regulated DEGs, 298 were found in the SUBA5 database and most of the encoded proteins were predicted to be active in the plasma membrane (23%), nucleus (21%) and cytosol (19%), while only 8% were targeted to plastids (**Fig. 9 C**).

**Figure 9.**
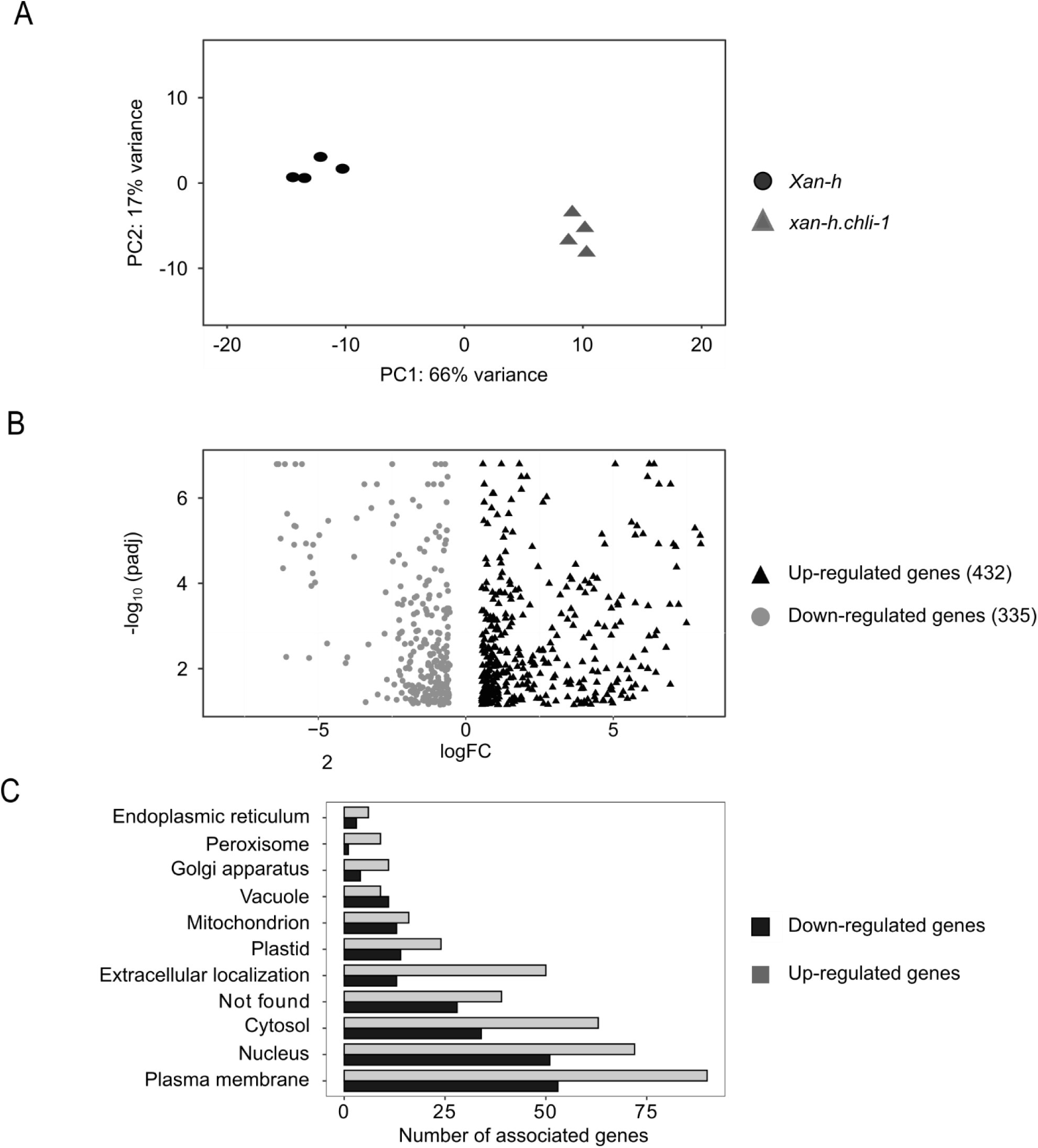
Comparative transcriptomic analyses of *xan-h.chli-1* and *Xan-h* leaves grown under greenhouse conditions. (A) Principal component analysis (PCA) of the four biological replicates for each genotype. (B) Volcano plot of the differentially expressed genes (DEGs) filtered by the log of fold change (logFC) and the adjusted p-value (padj). (C) Subcellular localization of DEGs based on information available in the SUBA5 database (https://suba.live/).

Also, in the case of the 177 down-regulated genes found in the SUBA5 database, the majority of the encoded proteins were localized to the plasma membrane (27%), the nucleus (24%) or the cytosol (16%), whereas only 7% of the genes encoded plastid proteins, further confirming the limited impact of the *xan-h.chli-1* mutation on chloroplast functionality.

In light of the localization of the Mg-chelatase enzyme and the pale green phenotype of mutant plants, the chloroplast-related DEGs were analysed in detail. Twenty-three up-regulated nuclear genes were predicted or reported to encode proteins active in the chloroplast (**Table 2**). These included the *ATNTH1* gene encoding a DNA glycosylase-lyase involved in base excision repair of oxidative DNA damages and an M-type 4 thioredoxin with a role in the oxidative stress response. Genes coding for proteins with a role in jasmonic-acid-mediated stress responses, such as lipoxygenae 2 (*LOX2*), the lipase *DALL4*, and the allene oxidase synthase (*AOS*), and in drought and heat-stress responses, as in the case of *Heat Shock Protein 21* (*HSP21*) and *TRR14,* were also upregulated (see **Table 2**). In addition, several upregulated genes are reported to play a role during the early stages of chloroplast and seedling development, including early light inducible protein 1 (*ELIP1*), plastid transcriptionally active chromosome 18 (*pTAC18*), raspberry 3 (*RSY3*), and arogenate dehydratase 6 (*ADT6*). Furthermore, the plastid type I signal peptidase 1 (*PLSP1*) and the plastid type I signal peptidase 2B (*PLSP2B*), which remove signal sequences from proteins translocated into the thylakoid lumen, were also upregulated.

**Table 2.** A subset of nuclear genes that code for proteins located in the chloroplast and are differentially expressed in *xan-h.chli-1* and *Xan-h* leaves under greenhouse conditions.

| Gene code<br>(HORVU.MOREX.r3) | Gene name | Biological process/function | LogFC | Ref |
| --- | --- | --- | --- | --- |
| Up-regulated genes in *xan-h.chli-1* vs *Xan-h* |  |  |  |  |
| 4HG0343330 | DNA glycosylase<br>superfamily protein<br>(ATNTH1) | Repair of oxidative DNA damages | 7,59 | (Gutman and Niyogi 2009) |
| 6HG0540480 | Lipoxygenase 2<br>(LOX2) | Jasmonic Acid-mediated stress<br>response | 6,94 | (Kaur et al. 2023) |
| 7HG0639230 | Lipoxygenase 2<br>(LOX2) | Jasmonic Acid-mediated stress<br>response | 6,12 | (Kaur et al. 2023) |
| 4HG0392440 | Chaperon DnaJ-domain superfamily<br>(DJC24) | Recruit Hsp70 to perform tissue<br>specific function | 2,78 | (Chiu et al. 2013) |
| 5HG0489830 | Heat Shock Protein<br>21 (HSP21) | Required for seedling and chloroplast<br>development under heat stress | 2,05 | (Sedaghatmehr,<br>Mueller-Roeber,<br>and Balazadeh<br>2016) |
| 5HG0420500 | Lipoxygenase 2<br>(LOX2) | Jasmonic Acid-mediated stress<br>response | 1,58 | (Chiu et al. 2013) |
| 3HG0323270 | Alpha/beta-<br>hydrolases<br>superfamily<br>(DALL4) | Jasmonic acid biosynthesis | 1,51 | (Rudus et al. 2014) |
| 4HG0394970 | Allene oxide<br>synthase (CYP74A) | Jasmonic acid biosynthesis | 1,35 | (Oshita et al. 2023) |
| 2HG0097390 | Cystathionine beta-<br>synthase family<br>protein (CBSX1) | Stabilize cellular redox homeostasis<br>and modulate plant development via<br>regulation of Thioredoxin systems | 1,15 | (Ok, Yoo, and Shin<br>2012) |
| 7HG0738610 | Thylakoid<br>processing<br>peptidase (TTP1) | Thylakoidal processing peptidase that<br>removes signal sequences from<br>proteins transported into the thylakoid<br>lumen | 1,12 | (Hsu et al. 2011) |
| 7HG0749460 | Thioredoxin M-type<br>4 (TRXM4) | Prokaryotic thioredoxin involved in<br>the response to oxidative stress and<br>control of cyclic electron transport<br>and Calvin cycle. | 0,86 | (Okegawa and<br>Motohashi 2015) |
| 5HG0482040 | Chlorophyll A-B<br>binding family<br>protein (ELIP1) | Prevent excess accumulation of free<br>chlorophyll by inhibiting the entire<br>chlorophyll biosynthesis pathway and<br>hence prevent photooxidative stress. | 0,76 | (Rizza et al. 2011) |
| 1HG0053910 | Plastid<br>Transcriptionally<br>Active Chromosome<br>18 (PTAC18) | Associated with the PEP and<br>chloroplast transcription activity and<br>chloroplast biogenesis | 0,75 | (Pfalz et al. 2006) |
| 6HG0595460 | RmlC-like cupins<br>superfamily protein<br>(TRR14) | Overexpression leads to trehalose<br>resistance, drought and stress<br>tolerance. *trr14* mutants have reduced<br>seed germination, root length,<br>survival rate and chlorophyll content<br>under stress conditions. | 0,71 | (Aghdasi, Fazli, and<br>Bagherieh 2012) |
| 2HG0165450 | *Arogenate oxygenase 6 (ADT6)* | A plastid-localized arogenate dehydratase involved in phenylalanine biosynthesis. | 0,69 | (Apuya et al. 2002) |
| 6HG0576310 | *Raspberry 3 (RSY3)* | chloroplast biogenesis and seedling development | 0,60 | (Apuya et al. 2002) |
| Down-regulated genes in *xan-h.chli-1* vs *Xan-h* |  |  |  |  |
| 7HG0739190 | *Cryptochrome 3 (CRY3)* | Bind flavin adenine dinucleotide and DNA. It is likely to act as photoreceptor | -0,60 | (Göbel et al. 2017) |
| 7HG0738860 | *Peptidyl-prolyl cis-trans isomerase (FKBP13)* | FKBP13 is located in chloroplast thylakoid lumen and possess peptidyl-prolyl cis/trans isomerase (PPIase) activity. | -0,68 | (Ingelsson et al., 2009) |
| 7HG0689310 | *Late embryogenesis abundant hydroxyproline-rich glycoprotein family (LEA)* | Function in normal plant growth and development, and in protecting cells from abiotic stress | -0,81 | (Hong-Bo, Zong-Suo and Ming-An, 2005) |
| 5HG0533290 | *ACYL ACTIVATING ENZYME 16 (AAE16)* | Fatty acid biosynthetic process | -1.06 | (Tjellström et al. 2013) |
| 4HG0418700 | *RUBISCO ASSEMBLY FACTOR 2 (RAF2)* | Protein involved in Rubisco assembly that also mediates Absciscic acid-dependent stress response. | -1.28 | (Fristedt et al. 2018) |
| 7HG0712530 | *Alpha amylase family protein (BE1)* | Putative glycoside hydrolase localized in plastids, plays crucial roles during embryogenesis and carbohydrate metabolism | -4,01 | (Wang et al. 2010) |
| 3HG0228020 | *Sulfoquinovosyl diacylglycerol 2 (SQD2)* | Involved in sulfolipid biosynthesis | -4.07 | (Okazaki et al. 2013) |
| 7HG0689300 | *Thioredoxin family protein* | Change in response to H<sub>2</sub>O<sub>2</sub>. | -10.06 | (Peltier et al. 2004) |

The thirteen chloroplast-located down-regulated genes are mainly involved in protein folding and assembly, such as FK506-binding protein 13, a peptidyl-prolyl isomerase located in chloroplast thylakoid lumen which is considered to act as a protein folding catalyst, and *RAF2*, Rubisco Assembly Factor 2, in fatty acid and lipid biosynthesis, such Acyl Activating Enzyme 16 (*AAE16*) and UDP-sulfoquinovose:DAG sulfoquinovosyltransferase 2 (*SQD2*), respectively, and during early developmental stages, as in the case of *BE1*, a putative glycoside hydrolase that plays a vital role during embryogenesis and in carbohydrate metabolism, and Late Embryogenesis Abundant (*LEA*) hydroxyproline-rich glycoprotein which is thought to function in plant development and growth. Strikingly, none of the nuclear genes encoding subunits of the thylakoid photosynthetic apparatus were among those differentially regulated by the *xan-h.chli-1* mutant allele.

### The *xan-h.chli-1* mutant line is characterised by reduced daily transpiration rate

To investigate the growth advantages associated with the pale green leaf phenotype, *xan-h.chli-1* and *Xan-h* plants were grown in pots in Plantarray, a functional phenotyping platform (FPP), in a semi-controlled environmental greenhouse, with the aim of detecting small changes in specific physiological processes under both optimal and limiting watering regimes (Lupo and Moshelion 2024). Plant biomass and water flux measurements performed throughout the entire plant life cycle, from January 19^th^ to March 2^nd^, 2023, allowed for calculations of transpiration and biomass gain of each plant (Appiah et al. 2023). Plants were initially grown under well-watered conditions for 23 days, followed by an 18-day period of drought-stress, and then returned to standard conditions until harvesting. Under well-watered conditions, the daily transpiration rate normalized to plant fresh weight was significantly lower in *xan-h.chli-1* plants (from 10 to 55% reduction) than in the *Xan-h* control, as shown in **Figure 10 A**, where data collected during 14 representative days are shown. Similar differences were observed during the drought stress period (**Fig. 10 B)**, where *xan-h.chli-1* plants showed a stable reduction in the daily transpiration rate, i.e. around 40-50% less than the *Xan-h* control. Strikingly, towards the end of the drought-stress period, when plants were under severe water deficiency, *Xan-h* daily transpiration rate decreased severely reaching values significantly lower than those recorded for *xan-h.chli-1*, most probably as a consequence of the fact that *xan-h.chli-1* plants were better able to tolerate the drought stress (**Fig. 10 B)**. However, this general reduction in transpiration rate comes at the expense of total biomass accumulation (*xan-h.chli-1* 16.52 ± 4.42 gr *vs. Xan-h* 25.34 ± 2.57 gr; **Fig. 10 C**) and water use efficiency (WUE), i.e. biomass gain per ml of transpired water (*xan-h.chli-1* 0.00364 ± 0.00064 gr ml^-1^ *vs Xan-h* 0.00446 ± 0.00036 gr ml^-1^; **Fig. 10 D**)

**Figure 10.**
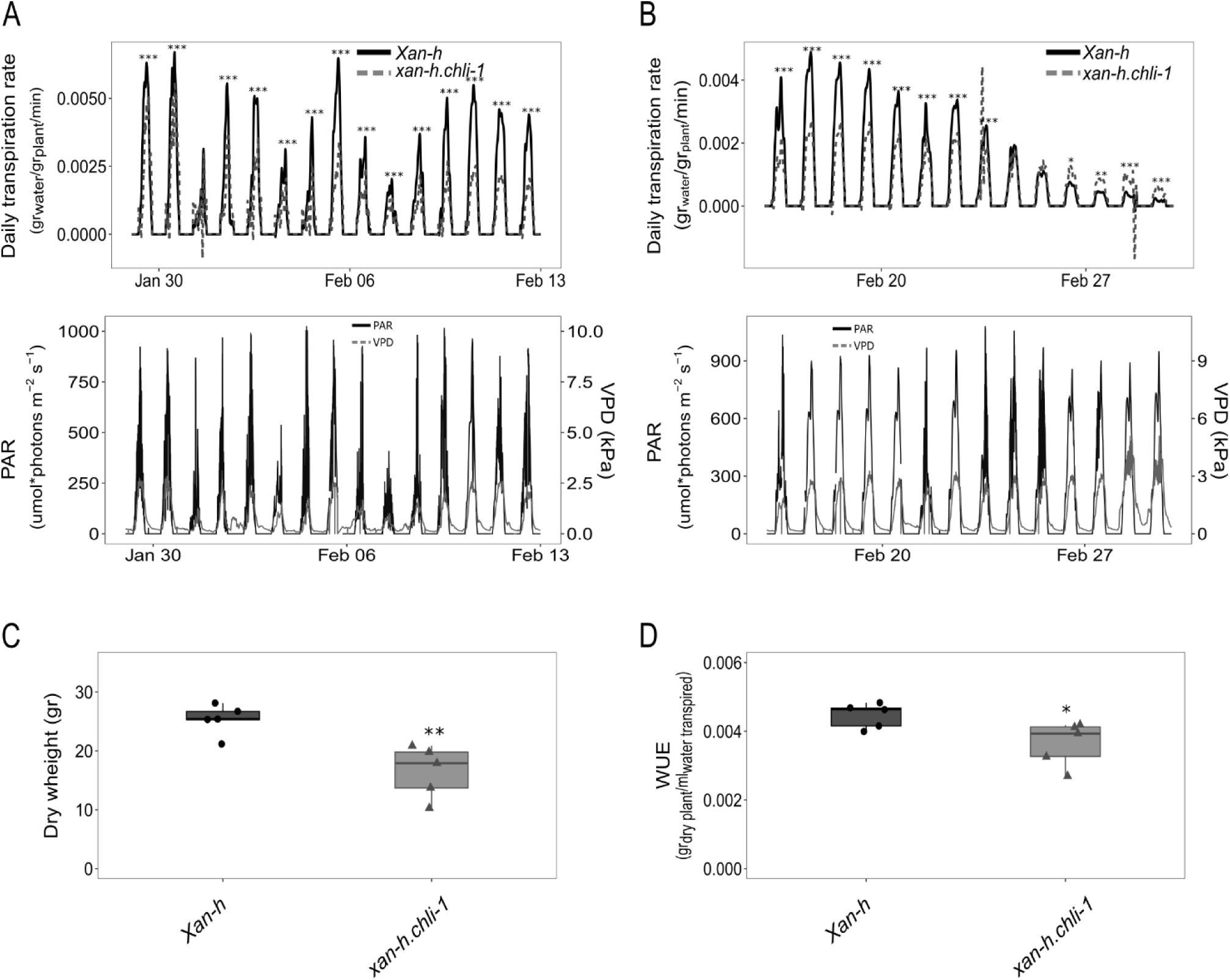
Relative performance of *Xan-h* and *xan-h.chli-1* plants grown under optimal and drought-stress conditions, as estimated by the FPP phenotyping platform. (A) Upper panel: Daily transpiration rate normalized to plant fresh weight (g water/g plant/min) as evaluated for 14 representative days under well-watered conditions during daylight exposure from 6.00 am to 18.00 pm. To avoid overloading the Figure, data obtained during the night period are not shown. Lower panel: Photosynthetic active radiation intensities (PAR) and vapour pressure deficit (VPD) measured by a weather station for the 14 representative days under well-watered conditions. Panel B and C depicts the corresponding data for plants grown for 14 representative days under drought-stress conditions, automatically maintained through the feedback-controlled irrigation system, during daylight exposure from 6.00 am to 18.00 pm. In all cases, the significance of the data was estimated using Student’s t-test (*** P < 0.001, ** P < 0.01, * P < 0.05). (C) Plant dry weight (g) at the end of the experiment, i.e. upon completion of plant life cycle. The plant material was dried at 60°C for 72 h. The significance of the observed differences was evaluated with Students t-test (** P < 0.01). (D) Water use efficiency (WUE) (g dry plant/ml water transpired) was measured by the total weight of dry plants at the end of the life cycle, normalized to total water transpired. Student’s t-test was performed to estimate the significance of the observed differences (* P < 0.05).

## Discussion

Our research interests focus on testing in crops the potential of new biotechnological strategies that have been shown to increase photosynthetic efficiency and yield in model organisms under controlled conditions (Burgess and Cardoso 2023; Long, Marshall-Colon, and Zhu 2015). Barley lends itself very well to this approach owing to the availability of large collections of natural and induced genetic diversity, complete genome sequences, protocols for genetic transformation and gene editing (Rotasperti et al. 2020), and proven, worldwide economic importance. Among the proposed strategies, we have focused our attention on the manipulation of leaf pigment content, which has been reported to enhance light use efficiency in high-density monocultures (Kirst et al. 2017). Many genetic targets are available for the alteration of leaf chlorophyll levels, as reviewed in Cutolo et al., 2023. Recently, we reported on the barley mutant *happy under the sun 1* (*hus1*), which is characterised by a 50% reduction in the chlorophyll content of leaves, owing to a premature stop codon in the *HvcpSRP43* gene that codes for the 43-kDa chloroplast Signal Recognition Particle (Rotasperti et al. 2022). The *Hv*cpSRP43 protein is responsible for the insertion of photosystem antenna proteins into the thylakoid membranes, and its truncation results in a decrease in antenna size. However, when sown at standard density under field conditions, the yield of *hus1* plants is comparable to that of the wild type, and the reduction in chlorophyll leaf content is well tolerated under conditions that are typical of cultivated fields. In the present work, we have characterized a novel chlorophyll-deficient mutant in barley. This pale-green mutant, *xan-h.chli-1,* is due to a missense mutation (R298K) in a highly conserved residue of the *Hv*CHLI protein – the smallest subunit of the Mg-chelatase enzyme, which catalyses the first unique step in chlorophyll biosynthesis (Lundqvist et al. 2010; 2013). Interestingly, while all of the mutants previously described at the *Xan-h* locus in barley (*xan-h.38*, *xan-h.56*, *xan-h.57* and *xan-h.clo125*, *xan-h.clo157*, *xan-h-clo161*) show a seedling-lethal phenotype (Hansson et al. 1999; Braumann, Stein, and Hansson 2014), the *xan-h.chli-1* line is viable (**Fig. 1, 3, 5**). This unique phenotype is due to the fact that the R298K missense mutation does not dramatically affect the accumulation of *Hv*CHLI, nor its ability to form homodimers, but rather results in a drastic reduction of the Mg-chelatase activity, as experimentally verified *in vitro* (see **Fig. 7**). The fact that the *xan-h.clo125*, *xan-h.clo157* and *xan-h.clo161* variants do not form homodimers in yeast two-hybrid assays (**Fig. 7**), while the corresponding variants of the *Rhodobacter capsulatus bchI* gene oligomerize on a gel-filtration column in the presence of ATP (Hansson et al. 2002), may be ascribed to the relatively low homology (49% identity) between their amino-acid sequences (**Fig. S1**).

The introgression of the *xan-h.chli-1* mutant allele into the lethal *Atchli1/Atchli1* mutant background (Huang and Li 2009) fully restored plant viability and reverted the albino *Atchli1/Atchli1* phenotype to a milder pale-green leaf colour, similar to those of the Arabidopsis *cs*/*cs* (Kobayashi et al. 2008) and barley *xan-h.chli-1* mutant phenotypes (see **Fig. 4**). Furthermore, the *xan-h.chli-1* allele also attenuated the lethal *xan-h.56* phenotype, as the biallelic mutant *xan-h.56/xan-h.chli-1* shows a pale green leaf phenotype and its photosynthetic efficiency is comparable to that of *xan-h.chli-1* leaves (see **Fig. 3**). Since the homozygous *xan-h.56* mutant does not accumulate the *Hv*CHLI protein (Braumann, Stein, and Hansson 2014), the pale-green phenotype of the heterozygote is attributable to the *xan-h.chli-1* allele alone. Conversely, the *xan-h.clo161* barley mutant showed a reduction in *Hv*CHLI accumulation relative to the wild-type, and the semi-dominant nature of this mutation indicates that the protein encoded by the *xan-h.clo161* allele has detrimental effects on the assembly and activity of the *Hv*CHLI hexamer (Hansson et al. 2002). This accounts for the accumulation of both protein variants in the barley biallelic mutant *xan-h.clo161*/*xan-h.chli-1*, in which the interaction of the two variants within the *Hv*CHLI hexamer most probably results in a non-functional Mg-chelatase and a lethal *Chlorina*-like phenotype (**Fig. 3**). Our findings are also in agreement with the localization of the R298 residue, which is predicted to reside in the ATP-binding pocket and to interact directly with the ATP molecule. The model displays conformational similarity to the recently published CHLI hexamer structure from *Synechocystis* sp. PCC 6803 (PDB ID 6L8D, Gao et al., 2020; **Fig. 8** and **Fig. S3**), which exhibits 73% sequence identity with WT *HvCHLI* (**Fig. S1).** In particular, the altered interactions caused by the R298K substitution in the ATP-binding pocket suggest that the R298 residue is involved in either ATP binding and/or ATP hydrolysis.

As expected, pale green *xan-h.chli-1* leaves showed a reduced content of both Chls *a* and *b* and an increase in the Chl*a*/Chl*b* ratio when compared to control plants (**Table 1**). Furthermore, comparable reductions were observed in the accumulation of carotenoids, including β-carotene which is preferentially associated with the PSI and PSII cores. This indicates that – unlike the alteration of antenna protein biogenesis in *hus1* mutant – the impairment of chlorophyll biosynthesis leads to a general destabilization of the entire photosynthetic apparatus, as is confirmed by the reduced accumulation of antenna proteins and photosystem core subunits observed by immunoblots (**Fig. 6**). In addition, the reorganization of electron transport in the thylakoids of *xan-h.chli-1* leaves appears to take place at the post-transcriptional level, as transcriptomic analysis revealed that none of the nuclear genes encoding photosynthesis-associated proteins were affected in the mutant (**Fig. 9**, **Table 2**).

Intriguingly, the reduction of antenna size in *xan-h.chli-1* leaves, together with the decline in photosystem core proteins, decreased the efficiency of photosynthesis only under low light intensities, whereas photosynthetic performance was enhanced relative to WT under high light levels (**Fig. 5**), as already reported in the case of *hus1* plants (Rotasperti et al. 2022). This is most probably due to the reduction in thylakoid excitation pressure in *xan-h.chli-1* leaves exposed to high-light intensities, as indicated by the lower values of Y(NPQ) and Y(ND) parameters (**Fig. 5**). Moreover, *xan-h.chli-1* plants are characterised by a significantly lower transpiration rate, at the expense of total biomass accumulation and WUE (**Fig. 10**). This behaviour, combined with the reduction in Chl*b* leaf content, resembles the high-risk drought escape strategy adopted by certain wild barley accessions (*Hordeum vulgare* spp. *spontaneum*) that are adapted to stable and very dry environments where fitness (i.e. reproductive output and the quality of offspring) prevails over the achievement of the full production potential (Galkin et al. 2018). This finding supports the notion that pale-green leaves may have beneficial effects in harsh environments, as in the case of certain Syrian barley landraces that grow under arid climatic conditions and are characterized by a pale green phenotype (Watanabe and Nakada 1999; Tardy, Créach, and Havaux 1998).

Overall, while crop breeding has led to the development of high-yielding cultivars, progress toward the development of crops that tolerate abiotic stresses has been very slow. Thus, the need to reduce the ‘yield gap’ and improve yields under a variety of stress conditions is of strategic importance for future food security (Sadras and Richards 2014; Cattivelli et al. 2008; Araus et al. 2002). In this context, the *xan-h.chli-1* mutant allele and its pale green phenotype have a significant potential for application in breeding programs that deserves to be investigated. These traits include a more equal distribution of light under high-density field conditions and potential benefits for net photosynthetic efficiency across the entire canopy, together with the possible benefits under drought condition and in the efficiency of nitrogen use (Walker et al. 2018; Sakowska et al. 2018). To this end, the introgression of the *xan-h.chli-1* allele into elite barley cultivars, and collaboration with plant breeders, agronomists and crop physiologists to select the most appropriate yield-testing protocols, definition of growing plant densities and standard parameters to define yields are needed. Furthermore, *xan-h.chli-1* seedlings do not show the *genomes uncoupled* phenotype in the presence of Norflurazon, unlike seedlings that carry lethal allelic variants (Fig. S4; Gadjieva et al. 2005). This is advantageous, since retrograde signalling plays a crucial role in the adaptation of plants to changing environments, and several *gun* mutants show impaired responses and heightened sensitivity to abiotic challenges (Song, Chen, and Larkin 2018; Marino et al. 2019). Finally, in the medium term, the knowledge gained could be transferred to other cereals, including wheat, given the high degree of conservation of the chlorophyll biosynthetic pathway and the photosynthetic machinery in higher plants.

## Materials and Methods

### Nucleotide and amino acid sequence analysis

Amino-acid and genome sequences of *Xan-h* (*HORVU.MOREX.r3.7HG0738240*), *AtCHLI1* (*At4g18480*) and *AtCHLI2* (*At5g45930*) were obtained from the ENSAMBLE-Plant database (plants.ensembl.org/index.html). Multiple sequence alignments were obtained locally with Muscle v5 (drive5.com/muscle5/) (Edgar 2022). Subcellular localization and chloroplast transit peptide (cTP) predictions were identified by TargetP (services.healthtech.dtu.dk/services/TargetP-2.0/).

### Plant material and growth conditions

Barley (*Hordeum vulgare*) plants were cultivated on acid soil (Vigor plant-growth medium, based on Irish and Baltic peats, pH 6.0) supplemented with Osmocote fertilizer under controlled greenhouse conditions (250 µmol photons m ^-2^ s ^-1^ for 16 h and 8 h dark). Temperatures were set to 20°C during the day and 16°C at night, with a relative humidity of 30%. The *TM2490* line was identified among the M_4_ generation of the chemically mutagenized TILLMore population (Talamè et al. 2008), which is derived from the ‘Morex’ cultivar background. The F_2_ segregating population, generated for mapping purposes, was obtained by manually crossing the *TM2490* line with the cv. Barke. The *xan-h.chli-1* line was isolated from an F_2_ population obtained by backcrossing *TM2490* with the barley *cv.* Morex (BC_2_F_2_). Arabidopsis Columbia-0 (Col-0) and mutant lines were grown on soil in a climate chamber (150 µmol photons m^-2^s ^-1^ for 16 h, and 8 h dark) at 22°C. The *Atchli1/Atchli1* T-DNA insertion mutant (*SAIL_230_D11*) was identified by searching the T-DNA Express database (signal.salk.edu/cgi-bin/tdnaexpress), while the homozygous line *cs/cs* was provided by Professor Tatsuru Masuda (Kobayashi et al. 2008). The transgenic Arabidopsis lines *Atchli1/Atchli1 + 35S::Xan-h* and *Atchli1/Atchli1 + 35S::xan-h.chli-1* were generated by *Agrobacterium-*mediated transformation of the heterozygous *Atchli1/AtCHLI1* mutant line with either the wild-type *Xan-h* or the mutant *xan-h.chli-1* coding sequence from barley, respectively, using the *pB2GW7* plasmid (VIB-UGhent for Plant Systems Biology). Primers used for mutant isolation and cloning procedures are listed in **Table S1**.

### The phenotyping platform

The functional-phenotyping platform Plantarray (FPP; PlantDitech Ltd.; Yavne, Israel) was used to monitor plant growth and water balance. The system monitored water transpiration and biomass gain of each plant throughout the growing period, while detecting environmental conditions every 3 min. A feedback irrigation system was used to ensure standardized drought treatment. All plants were exposed to comparable drought stress by taking each plant’s transpiration rate into account, as previously described (Dalal et al. 2020). Plants were grown on the Plantarray system for 43 days from January 19 to March 2, 2023.During the pre-drought phase of 24 days, all plants were well-watered at pot capacity through nocturnal irrigation. Drought conditions were progressively imposed from February 12 to March 2 (18 days) by gradually reducing the daily irrigation to 80%. After 10 more days on the Plantarray (recovery phase), plants were moved to the greenhouse and, upon harvest, total dry biomass weight was measured. To determine the dry weight, the plant material was dried at 60°C for 72 h (Appiah et al. 2023). Whole-plant transpiration rates were derived by multiplying the first derivative of the measured load-cell time series by −1. The transpiration was then normalized to the plant’s fresh weight. Water-use efficiency (WUE) was calculated as harvest product dry weight (g)/water transpired over the entire Plantarray growth period (ml). The accuracy of calculations was determined independently (Jaramillo Roman et al. 2021).

### RNAseq analyses for mapping and differential gene expression

For gene mapping analyses, RNA was isolated from 100 wild-type-like and 100 *TM2490*-like F_2_ plants (*TM2490* × *cv* Barke) and extracted as previously described (Verwoerd, Dekker, and Hoekema 1989). Independent RNA samples were bulked in equal ratio to generate two pools. RNA pools were subjected to poly-A capture and paired-end sequencing, producing approximately 100 million 2 × 150-bp read pairs per pool (47.97M and 48.89M 2 × 150 nt paired-end reads, for wild-type-like and *TM2490*-like pools, respectively). Reads were mapped to the *H. vulgare* Morex v3 genome sequence (Mascher et al. 2021). Coherent mapping was obtained for 84.5% and 86.1% of pairs for the wild-type-like and *TM2490*-like pools, respectively. Samtools (Danecek et al. 2021) was used to sort and index the resulting alignment files, and FreeBayes (v1.3.2) (Garrison and Marth 2012) was used to call variants, employing default parameters except for requiring a minimum mapping quality of 20 and a minimum base-call quality of 30. Custom Python scripts were employed to identify variants segregating between pools (allele frequency >0.1 in both pools and <0.9 in the wild-type-like pool). Variants were binned in 2-Mb intervals along the barley genome. Variants in this region (chr7H: 592500000..605500000) were analysed using the Ensembl-vep pipeline (McLaren et al. 2016).

For quantitative transcriptomic analyses, total RNA was isolated from 14-day-old leaf samples obtained from four biological replicates each of *Xan-h* and *xan-h.chli-1* plants. Differential gene expression analysis was performed in R using the DESeq2 package (Love, Huber, and Anders 2014). DEGs were filtered for log of fold change (logFC) >0.5 and an adjusted p-value (padj) <0.05. The ncbi-blast-2.14.0+ tool (ftp.ncbi.nlm.nih.gov/blast/executables/blast+/LATEST/) was used to perform identifier mapping on the barley genes (i.e., proteins in *A. thaliana* that appear to match the input protein sequences of *H. vulgare cv*. Morex with a given percentage of identity). The subcellular localization was predicted with SUBA5 (suba.live/index.html). Biological Process Gene Ontology (GO) term enrichment was performed with agriGO v2.0 (systemsbiology.cau.edu.cn/agriGOv2/specises_analysis.php?SpeciseID=1&latin=Arabidopsis_thal iana) (Tian et al. 2017). The raw RNASeq data have been deposited in the NCBI data repository (https://submit.ncbi.nlm.nih.gov/subs/bioproject/) under the bioproject identifier PRJNA1052990.

### Assay for genomes uncoupled

Barley and Arabidopsis seeds were surface-sterilized and grown for 6 days (100 μmol photons m^−2^s^−1^ on a 16 h/8 h light/dark cycle) on Murashige and Skoog medium (Duchefa, Haarlem, The Netherlands), supplemented with 2% (w/v) sucrose and 1.5% (w/v) Phyto-Agar (Duchefa). In order to discriminate the homozygous *xan-h.56* mutants, the barley seedlings were then transferred onto MS media supplemented with 5 µM NF, while *Arabidopsis* seeds were grown directly on NF-supplemented media. RNA was extracted from the seedlings, and cDNA was obtained using the iScript™ gDNA Clear cDNA Synthesis Kit (Bio-Rad). The *genomes uncoupled* (*gun*) phenotype was identified by monitoring the expression of *Rbcs* and *Lhcb3* genes in *Xan-h* and *xan-h.chli-1*, together with Arabidopsis Col-0, *cs/cs, 35S::Xan-h* and *35S::xan-h.chli-1* lines, using RT-qPCR. Primers are listed in **Table S1**. Arabidopsis *gun5* and barley *xan-h.56* mutants were used as positive controls for the *genomes uncoupled* phenotype.

### Yeast two-hybrid assay

Coding sequences for *Xan-h*, *xan-h.chli-1*, *xan-h.clo125*, *xan-h.clo157* and *xan-h.clo161*, devoid of cTPs, were cloned into *pGBKT7-GW* and *pGADT7-GW* (Takara Bio) vectors through Gateway cloning. Primer sequences are listed in **Table S1**. Yeast strains Y187 and AH109 were transformed according to the Clontech User’s Manual (PT1172-1) with the vectors *pGBKT7* and *pGADT7*, respectively, harbouring the WT *Xan-h* and the mutant variants. Each Y187 strain was mated with the respective AH109 strain and plated on synthetic drop-out (SD) medium lacking tryptophan (-W) and leucine (-L), to select for positive diploids. To test *Xan-h* and mutant variants homodimerization, overnight liquid cultures were normalized to OD 0.5 and plated on selective media lacking histidine (-W-L-H) and histidine and adenine (-W-L-H-A). The growth of yeast culture dilutions was observed after three days.

### Immunoblot analyses

Thylakoids and total protein extracts were prepared from equal amounts of barley leaves (fresh weight) collected from 2-week-old seedlings as described previously (Bassi and Simpson, 1987). Protein extracts were fractionated on denaturing 12% (w/v) acrylamide Tris-glycine SDS-PAGE (Schägger and von Jagow, 1987) and transferred to polyvinylidene-difluoride (PVDF) membranes (Ihnatowicz et al. 2004). Three replicate filters were probed with specific antibodies. Signals were detected by enhanced chemiluminescence (GE Healthcare). Antibodies directed against Lhca1 (AS01 005), Lhca2 (AS01 006), Lhca3 (AS01 007), Lhcb1 (AS01 004), Lhcb2 (AS01 003), Lhcb3 (AS01 002), Lhcb4 (AS04 045), Lhcb5 (AS01 009), D1 (AS05084), D2 (AS06 146), CP43 (AS111787), CP47 (AS04 038), PsaA (AS06172), PsaD (AS09461), PetA (AS08 306), PetE (AS06 141), PsbO (AS05092), PsbQ (AS06 142-16), PsbR (AS05 059), PsbS (AS09533), H3 (AS10710) were obtained from Agrisera (Vännäs, Sweden). Antibodies were raised against *Hv*CHLI, *Hv*CHLD, *Hv*CHLH as previously described (Lake et al. 2004). The *Hv*GUN4-specific antibody was kindly provided by Professor Mats Hansson (Lund University, Sweden). Three biological replicates were analysed for each SDS-PAGE and immunoblot.

### Mg-chelatase activity assay

*In-vitro* Mg-chelatase activity assays were performed according to Hansson et al. (1999). WT and *xan-h.chli-1* seeds were sown in vermiculite and grown in the dark for 10 days. Etiolated seedlings were then homogenized in 0.4 M mannitol, 20 mM Tricine-NaOH pH 9 and 1 mM DTT. Intact chloroplasts were enriched by 15-min centrifugation at 3000 g and loaded onto a 40% (vol/vol) Percoll cushion in homogenization buffer. Gradients were centrifuged for 15 min at 13000 g. After washing steps in homogenization buffer, chloroplasts were resuspended in 200 μL of lysis buffer (20 mM Tricine-NaOH pH 9, 1 mM DTT, 1 mM PMSF). After a centrifugation step at 11000 g for 5 min, the recovered supernatants containing the Mg-chelatase subunits were adjusted to the same protein concentration. The enzymatic assay was carried out by adding 1 μL of the reaction cocktail (50 mM ATP, 250 mM creatine phosphate, 250 mM MgCl_2_, and 0.06 mM deuteroporphyrin). Reactions were stopped by adding 1 mL acetone/water/25% ammonia (80/20/1, vol/vol/vol) and 200 µL heptane was added in order to remove chlorophyll from samples. To measure the relative amount of Mg-deuteroporphyrin, the emission spectrum of the acetone phase was recorded from 550 to 600 nm using an excitation wavelength of 408 nm. Excitation and emission slits were set to 5 nm.

### Pigment extraction and quantification

Pigments from Arabidopsis and barley were extracted from fresh leaves with 90% acetone. To determine Chl*a* and Chl*b* concentrations, spectrophotometric measurements were carried out according to Porra, Thompson and Kriedemann (1989) and normalized relative to fresh leaf weight. Barley leaf-pigment content was also estimated by reversed-phase HPLC (Färber et al. 1997) normalised to fresh weight. Measurements were performed on five biological replicates for each genotype. Chlorophyll content was also measured *in vivo* at different development stages using the SPAD-502 chlorophyll meter (Konica-Minolta, Tokyo, Japan).

### Chlorophyll fluorescence measurements

*In-vivo* Chl*a* fluorescence was recorded on second barley leaves with a Dual PAM 100 (Walz, Effeltrich, Germany) according to Barbato et al., 2020. After 30 min of dark adaptation, leaves were exposed to increasing actinic light intensities (0-1287 μmol photons m^−2^s^−1^) and the following thylakoid electron-transport parameters were determined: the effective quantum yields of PSII [Y(II)] and PSI [Y(I)], the PSII quantum yield of non-regulated energy dissipation [Y(NO)], the PSII quantum yield of regulated energy dissipation [Y(NPQ)], the quantum yield of non-photochemical energy dissipation in PSI due to acceptor side limitation [Y(NA)], and the non-photochemical PSI quantum yield of donor-side limited heat dissipation [Y(ND)] parameters. An imaging Chl fluorometer (Imaging PAM; Walz) was used to measure Chl*a* fluorescence, and for *in-vivo* imaging. Dark-adapted plants were exposed to the blue measuring beam (1 Hz, intensity 4; F_0_) and a saturating light flash (intensity 10) was used to determine Fv/Fm values. A 5-min exposure to actinic light (56 μmol photons m^-2^s^-1^) was then used to calculate Y(II). The Handy PEA fluorometer (Hansatech Instruments Ltd., UK) was used to measure Fv/Fm values in barley plants grown under field conditions during the screening of the TILLMore population, and under greenhouse conditions during plant growth.

### Transmission electron microscopy (TEM)

TEM analyses were performed as described previously (Tadini et al. 2020). Portions (2 mm × 3 mm) of the second leaves of *Xan-h* and *xan-h.chli-1* barely plants were manually dissected and fixed under vacuum in 2.5% (w/v) glutaraldehyde and 0.1 M sodium cacodylate buffer. After washing with water several times, samples were counterstained with 0.5% uranyl acetate (w/v) overnight at 4°C. Tissues were then dehydrated in increasing concentrations of ethanol (70%, 80%, 90%, 100% v/v) and permeated twice with 100% (v/v) propylene oxide. Samples were gradually infiltrated first with a 1:2 mixture of Epon-Araldite and propylene oxide for 2 h, then with Epon-Araldite and propylene oxide (1:1) for 1 h and left in a 2:1 mixture of Epon-Araldite and propylene oxide overnight at room temperature. Epon-Araldite resin was prepared by mixing Embed-812, Araldite 502, dodecenylsuccinic anhydride (DDSA) and Epon Accelerator DMP-30 according to the manufacturer’s specifications. Ultra-thin sections of 70 nm were cut with a diamond knife (Ultra 45°, DIATOME) and collected on copper grids (G300-Cu, Electron Microscopy Sciences). Samples were observed by transmission electron microscopy (Talos L120C, Thermo Fisher Scientific) at 120 kV. Images were acquired with a digital camera (Ceta CMOS Camera, Thermo Fisher Scientific).

### *Hv*CHLI hexamer structure prediction

The homo-hexameric ring model of the barley *Hv*CHLI ATPase subunit of Mg-chelatase was generated using version 3 of Multimer from DeepMind Alphafold2 (Evans et al. 2022), allowing for template search with HH-suite (Steinegger et al. 2019). The structure underwent relaxation by the gradient-descent method using Amber (Hornak et al. 2006) force-field. Additional refinement was performed with Protein Preparation Wizard (Madhavi Sastry et al. 2013) from the Schrödinger Maestro suite, version 13.7.125, release 2023-3 (Maestro, Schrödinger, LLC, New York, NY, 2023). From the same suite, the module Residue Scanning and Mutation was used to replace R298 with lysine (K298), allowing for side-chain prediction with backbone sampling up to 2.5 Å from the mutation site, and Glide XP (Friesner et al. 2006) to perform the docking of ATP. The open-source software PyMOL (Schrödinger, LLC, The PyMOL Molecular Graphics System, Version 2.6) was used for visualisation of the molecular structures and rendering of the Figures. The WT model of the *Hv*CHLI subunit is available at https://www.modelarchive.org/doi/10.5452/ma-xoqwu, while the R298K model is available at https://www.modelarchive.org/doi/10.5452/ma-tvik6.

## Acknowledgements

This study was carried out within the Agritech National Research Center, spoke 1 and received funding from the European Union (Next-Generation EU: PIANO NAZIONALE DI RIPRESA E RESILIENZA (PNRR) – MISSIONE 4 COMPONENTE 2, INVESTIMENTO 1.4 – D.D. 1032 17/06/2022, CN00000022). The experiments performed using the functional-phenotyping platform Plantarray were supported by the project Plant-RED, funded by the Italian Ministry of Foreign Affairs and International Cooperation and the Ministry of Science and Technology of the State of Israel (Italy-Israel Joint Call for Proposals on Scientific and Technological Cooperation 2021). The Mg-chelatase activity assay performed in the lab of Mats Hansson was supported by the Horizon Europe programme under Grant Number 101082091 – BEST-CROP. We thank Tatsuru Masuda for providing the Arabidopsis *cs/cs* homozygous line and we are grateful to Paul Hardy for critical reading of the manuscript. The NoLimits platform at University of Milano is acknowledged for the TEM analysis. This manuscript reflects only the authors’ views and opinions, neither the European Union nor the European Commission can be considered responsible for them.

## Authors contributions

AP, LT, LR, LC, AlTo and PP designed the study. SS, LT, SR and PP took care of *TM2490* mutant isolation. AP, LR, MH and VT performed the molecular, biochemical and physiological characterization of the mutants described. LT, VT, CB and AnTa contributed to the yeast two-hybrid assay. AP and LR were responsible for the TEM images. LR, DH and AP took care of sequencing data analysis and identification of *xan-h.chli-1* mutation. AP, VT and PJ performed pigment analyses. AD and MM conducted the drought-stress tests. FB and CC took care of protein structure prediction and modeling. All authors helped draft the manuscript. PP coordinated the study and took care of the final version of the manuscript. All authors gave final approval for publication and agree to be held accountable for the work performed therein.

