## Supplementary Figures for "A missense mutation in the *Xan-h* gene encoding the Mg-chelatase subunit I leads to a viable pale green line with phenotypic features of potential interest for barley breeding programs"

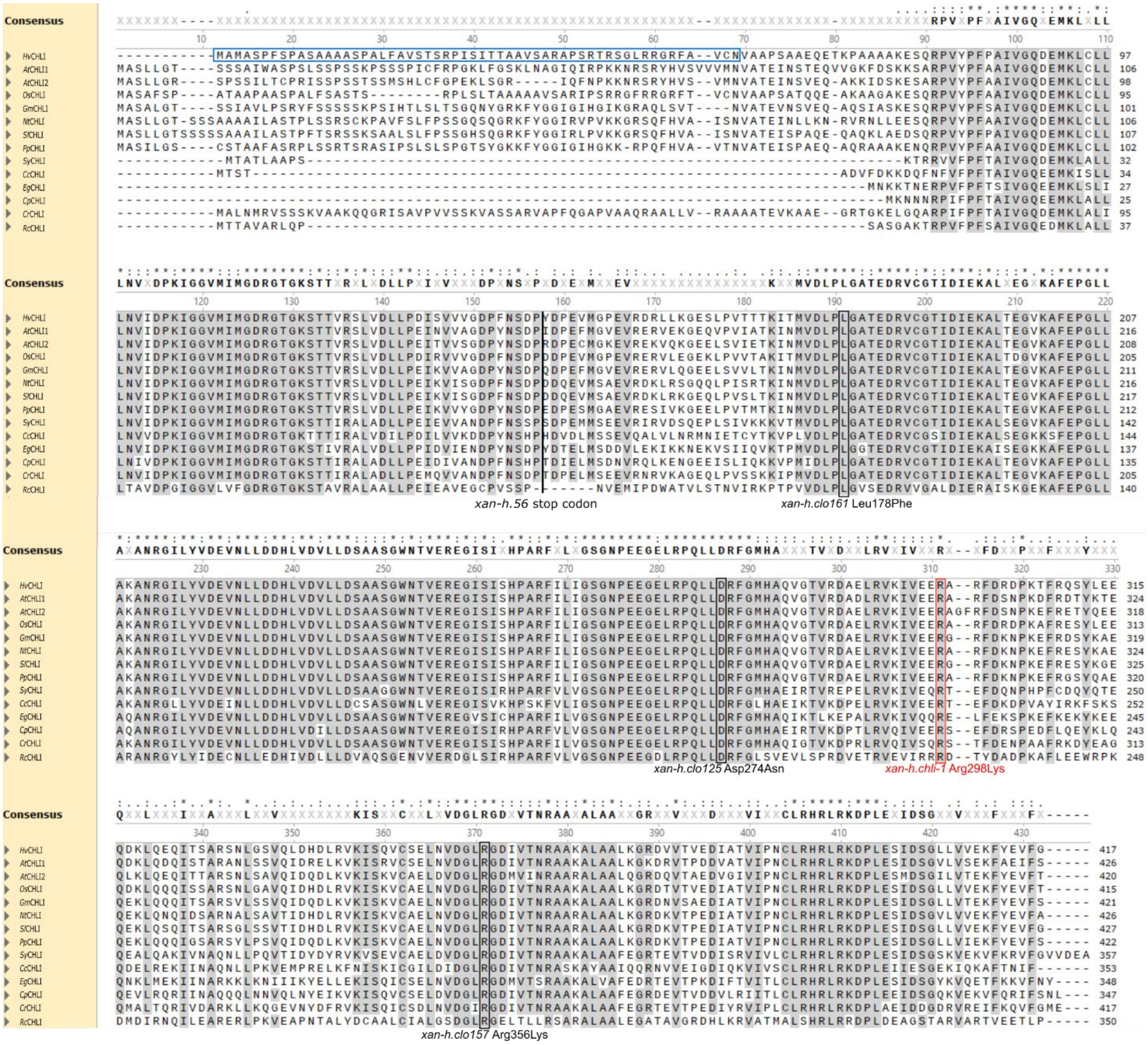


**Figure S1.** Alignment of CHLI sequences with Musclev5. The chloroplast transit peptide (cTP) of the *Hv*CHLI protein, as predicted by TargetP-2.0 (services.healthtech.dtu.dk/services/TargetP-2.0), is highlighted with a light-blue box. Amino-acid substitutions reported as SNPs in barley mutants are indicated. Amino-acid positions refer to the barley sequence, including the R298K missense mutation as main candidate for the *TM2490* pale green phenotype (in red). The degrees of identity between the *Hordeum vulgare* sequence and the analyzed sequences are the following: *Arabidopsis thaliana At*CHLI1 78%; *Arabidopsis thaliana At*CHLI2 81%; *Oryza sativa subsp. Japonica Os*CHLI 90%; *Glycine max Gm*CHLI 77%; *Nicotiana tabacum Nt*CHLI 76%; *Solanum lycopersicum Sl*CHLI 78%; *Prunus persica Pp*CHLI 78%; *Synechocystis sp.* (strain PCC 6803 / Kazusa) *Sy*CHLI 73%; *Cyanidium caldarium Cc*CHLI 62%; *Euglena gracilis Eg*CHLI 69%; *Cyanophora paradoxa Cp*CHLI 70%; *Chlamydomonas reinhardti Cr*CHLI 66%; *Rhodobacter capsulatus Rc*CHLI 49%.

Amino acids showing 100% conservation among the sequences considered are indicated by asterisks, while dots and colons indicate degrees of amino-acid conservation greater than 40% and 60%, respectively.

A

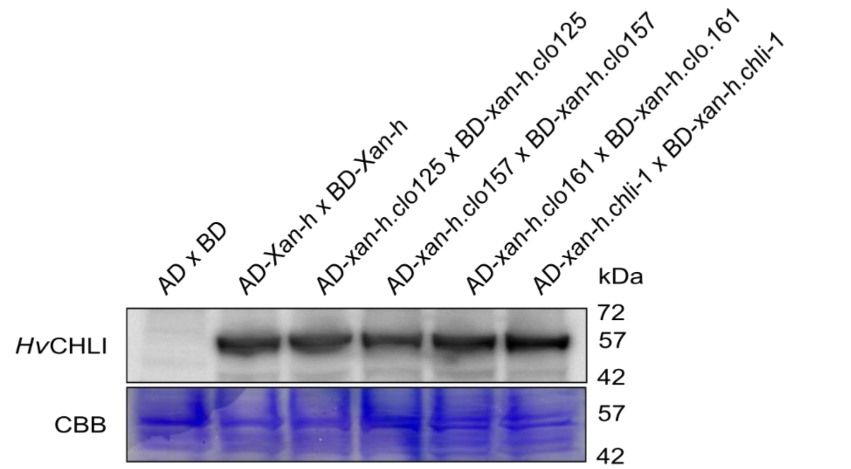


B

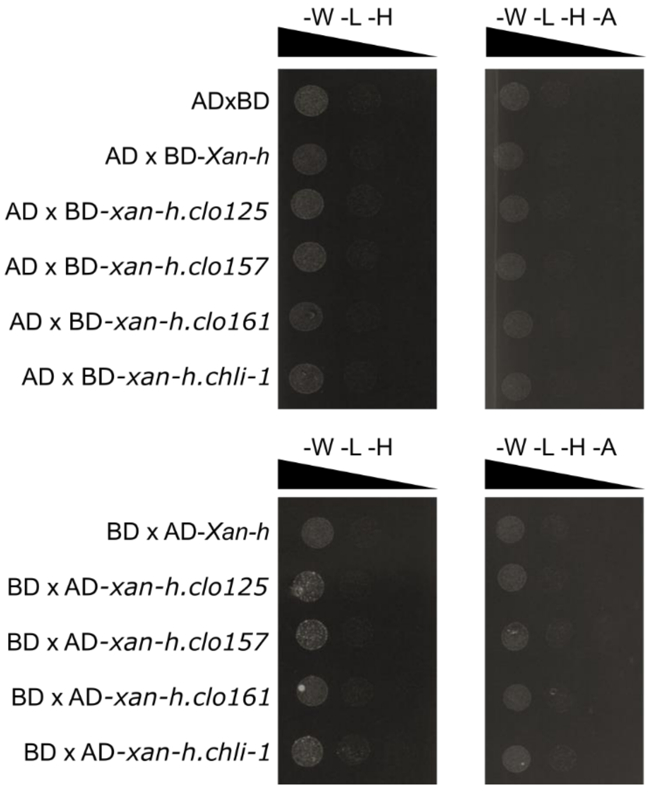


**Figure S2**. Western blots and negative controls to validate the yeast two-hybrid data. (A) Western blot analysis performed by using a *Hv*CHLI-specific antibody on diploid yeast cells (AH109xY187) to confirm the expression of Gal4AD and Gal4BD fusions to *Xan-h* and its allelic variants *xan-h.chli-1*, *xan-h.clo125*, *xan-h.clo157* and *xan-h.clo161* (MW between 53 and 57 kDa). Empty plasmids expressing Gal4AD and Gal4BD (AD x BD) were used as controls. CBB, Coomassie Brilliant Blue staining of a replica SDS-PAGE. (B) To exclude possible growth on selective media due to non-specific interaction between the different variants and the Gal4 domains used for the assay, each yeast strain expressing wild-type and mutant variants of *HvCHLI* was alternatively mated and tested for interaction against either the Gal4BD or the Gal4AD alone. No interaction between *Hv*CHLI and mutant variants with Gal4BD or Gal4AD could be detected, as shown by the lack of yeast growth on selective media, devoid of either Trp, Leu and His (-W -L -H) or Trp, Leu, His and Ade (-W -L -H -A).

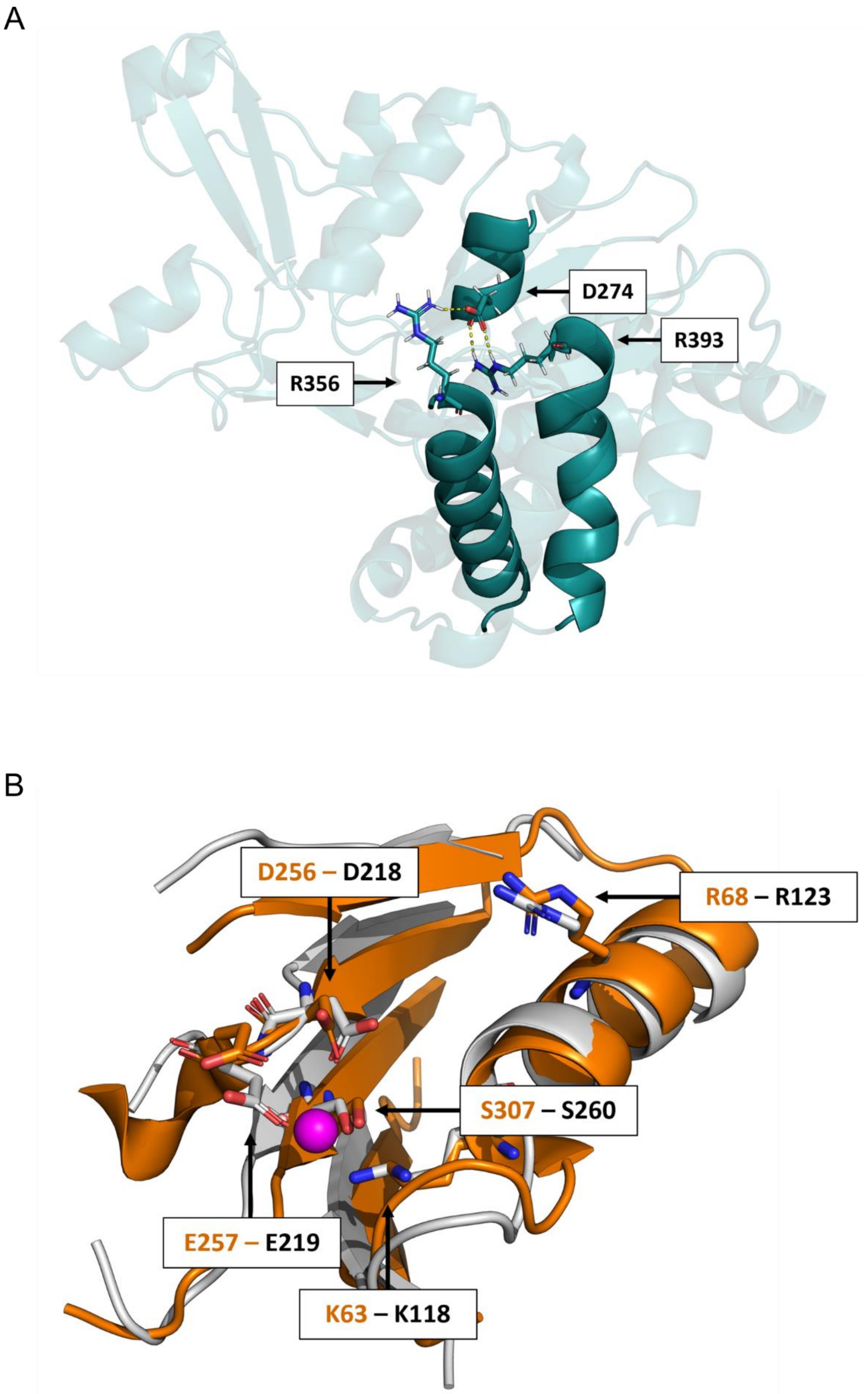


**Figure S3**. Details of specific properties of the *Hv*CHLI monomer. (A) Detailed view of the D274-R356-R393 interaction within the barley *Hv*CHLI monomer. The overall monomer structure is represented in transparent cyan cartoon, except for the alpha helices encompassing D274, R356 and R393, which are highlighted in solid cyan cartoon. D274, R356 and R393 are depicted as solid sticks and coloured: cyan for C atoms, blue for N, red for O, and white for H-. The hydrogen bond is represented as a dashed yellow line. (B) The ATP-binding domain from the protein Heat Shock Locus U [HSLU, PDB ID: 1DO0; (Bochtler et al. 2000)] superimposed on the barley *Hv*CHLI model. The HSLU domain is presented in solid orange cartoon, whereas the domain from the barley *Hv*CHLI model is in light grey. Selected conserved residues from HSLU are represented as solid sticks: C atoms are shown in orange, N atoms in blue, O atoms in red, and H atoms in white. Conserved residues of the barley model are depicted in the same colour scheme, except that C atoms are shown in light grey.

A B

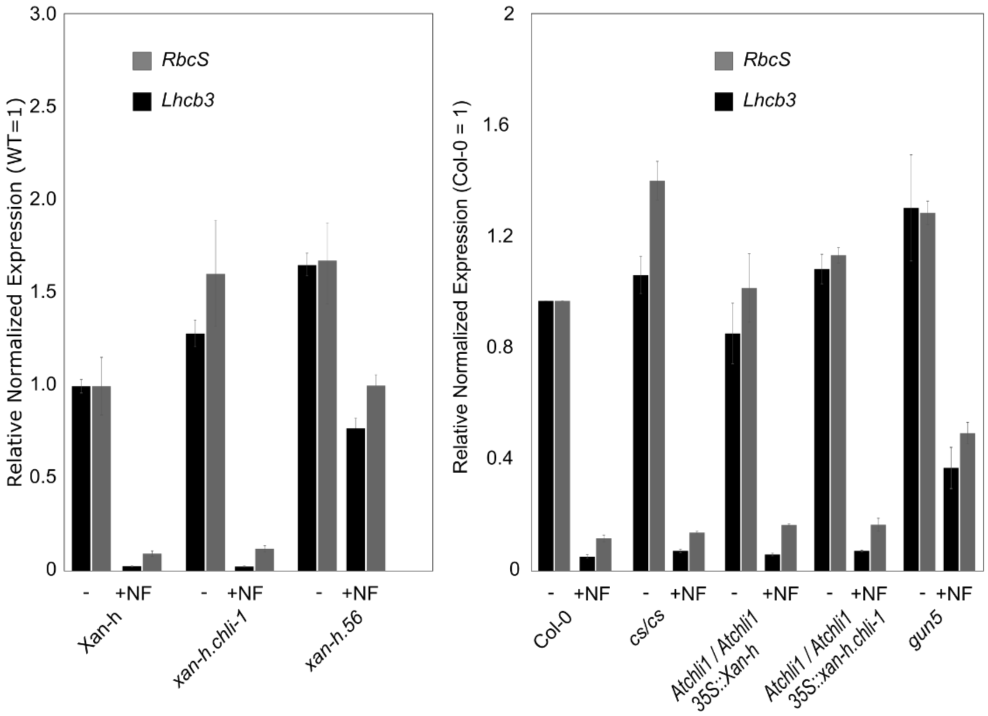


**Figure S4.** Assay for the *genomes uncoupled* (*gun*) phenotype in barley and Arabidopsis lines. (A) RT-qPCR expression analyses of the photosynthesis-associated nuclear genes *Rbcs* and *Lhcb3* were performed on barley *Xan-h*, *xan-h.chli-1* and *xan-h.56* lines grown for 6 days under sterile conditions, either in the absence of Norflurazon (NF) or on NF-supplemented medium (5 µM) for 4 days. (B) The expression of the same genes was also monitored in Arabidopsis Col-0, *cs*/*cs*, *Atchli1/Atchli1* + *35S::Xan-h* and *Atchli/Atchli* + *35S::xan-h.chli-1* mutant lines under the same conditions in presence or absence of 5 µM Norflurazon. The retrograde-signalling-defective mutant *gun5* (Arabidopsis) and *xan-h.56* (barley) were used as controls for the *gun* phenotype.
